# Distinct checkpoint and homolog biorientation pathways regulate meiosis I in *Drosophila* oocytes

**DOI:** 10.1101/2024.08.21.608908

**Authors:** Joanatta G. Shapiro, Neha Changela, Janet K. Jang, Jay N. Joshi, Kim S. McKim

**Author notes:** **Correspondence:** Kim S. McKim Corresponding author: Kim S. McKim 190 Frelinghuysen Road, Piscataway, NJ-08854.

## Abstract

Mitosis and meiosis have two mechanisms for regulating the accuracy of chromosome segregation: error correction and the spindle assembly checkpoint (SAC). We have investigated the function of several checkpoint proteins in meiosis I of *Drosophila* oocytes. Evidence of a SAC response by several of these proteins is found upon depolymerization of microtubules by colchicine. However, unattached kinetochores or errors in biorientation of homologous chromosomes does not induce a SAC response. Furthermore, the metaphase I arrest does not depend on SAC genes, suggesting the APC is inhibited even if the SAC is silenced. Two SAC proteins, ROD of the ROD-ZW10-Zwilch (RZZ) complex and MPS1, are also required for the biorientation of homologous chromosomes during meiosis I, suggesting an error correction function. Both proteins aid in preventing or correcting erroneous attachments and depend on SPC105R for localization to the kinetochore. We have defined a region of SPC105R, amino acids 123-473, that is required for ROD localization and biorientation of homologous chromosomes at meiosis I. Surprisingly, ROD removal, or “streaming”, is independent of the dynein adaptor Spindly and is not linked to the stabilization of end-on attachments. Instead, meiotic RZZ streaming appears to depend on cell cycle stage and may be regulated independently of kinetochore attachment or biorientation status. We also show that dynein adaptor Spindly is also required for biorientation at meiosis I, and surprisingly, the direction of RZZ streaming.

**Author Summary:** The Spindle Assembly Checkpoint (SAC) is known to delay cell cycle progression until chromosomes are properly attached to microtubules. Meiotic cells often have modified cell cycle phases, and natural arrest points such as metaphase I in *Drosophila*. We show that in *Drosophila* oocytes, the SAC is sensitive to loss of microtubules, but not sensitive to a variety of kinetochore attachment errors. Thus, the function of the SAC appears to be limited to monitoring oocyte spindle assembly, and not required for accurate chromosome segregation. However, two of the SAC genes, *rod* and *Mps1*, are required for the biorientation of homologous chromosomes during meiosis I, suggesting an error correction function. Rod is part of the RZZ complex and is notable for its property of streaming off the kinetochores. However, our results show that streaming off the kinetochore may not contribute to RZZ regulation of microtubule attachments, and only be associated with SAC function. Instead, the establishment of stable end-on attachments may occur while RZZ is still present at kinetochore. We suggest that RZZ interacts with multiple motors to promote bidirectional movement of kinetochores along microtubules, which allows chromosomes to find and attach to the correct pole.

## Introduction

Errors during meiotic chromosome segregation can result in inviable gametes leading to spontaneous abortions and infertility, or viable offspring with chromosomal abnormalities. Several mechanisms exist in dividing cells to prevent aneuploidy resulting from improper kinetochore-microtubule (KT-MT) attachments. The spindle assembly checkpoint (SAC) delays anaphase onset until attachments form between kinetochores and microtubules [1, 2]. Unattached kinetochores generate a signal involving SAC proteins such as MAD1, MAD2, BUB1, and BUB3 that have a global effect on all chromosomes and cell cycle progression. In contrast, error correction (EC) is specific to individual kinetochores whereupon interactions between microtubules and properly bi-oriented kinetochores are stabilized while those with improperly oriented kinetochores are weakened [3]. Multiple models have been proposed to explain how improperly oriented kinetochores are identified, involving both tension-dependent and -independent mechanisms, and how they are corrected, including phosphorylation of kinetochore proteins to destabilize attachments [4, 5].

Error rates are higher in human oocytes than in spermatocytes, which has been attributed to several causes [6–8]. One hypothesis is that the SAC is weak or inefficient [7, 9–11]. Indeed, Xenopus oocytes may not have a functional SAC [12, 13]. The observation that mammalian oocytes progress through anaphase with segregation errors may be a result of a weak checkpoint [14, 15]. The reason for a weak SAC in oocytes is not known, although possible contributing factors include the large size of oocytes or the lack of centrosomes [9, 16–19]. The SAC may also have other functions in oocytes, such as regulating the development of the oocytes and the timing of meiotic progression [10].[1, 2].

*Drosophila* oocytes are an excellent model for studying meiosis in females, but the SAC has not been examined. *Drosophila* oocytes have kinetochores that include the KMN complex (KNL1-MIS12-NDC80) [20]. Biorientation of homologous chromosomes involves types of kinetochore-microtubule attachments, lateral attachments that depend on KNL1/SPC105R and end-on attachments that depend on NDC80 [21, 22]. To understand the contributions of SAC and EC to biorientation in *Drosophila* oocytes, we characterized the SAC response during the first meiotic division, investigated how the SAC proteins are recruited to the kinetochores, and tested several SAC proteins for functions in meiotic EC. Our data suggest that the SAC is not sensitive to kinetochore attachment status and might be silenced prior to the metaphase I arrest. Furthermore, among seven SAC proteins, only two, ROD of the ROD-ZW10-Zwilch (RZZ) complex and MPS1, are required for biorientation of meiotic homologous chromosomes at metaphase I.

We examined the role of RZZ in *Drosophila* oocytes because it has not been studied before in meiosis. Previous studies in *C. elegans* embryos and *Drosophila* neuroblasts suggest that RZZ prevents errors by inhibiting premature stabilization of interactions between the ends of microtubules and NDC80 [23, 24]. RZZ also has the property of streaming off the kinetochore, which could alleviate the inhibition of end-on attachments [24–26]. We have shown that the RZZ complex is required for accurate homolog biorientation in meiosis I, which could indicate a role for the RZZ complex in delaying the formation of end-on attachments. We also show that Spindly, a dynein adaptor for RZZ, is also required for homolog biorientation in meiosis I. However, RZZ streaming in oocytes is not required for, nor does it depend on, the formation of end-on attachments. We propose that RZZ streams in response to the silencing of the SAC prior to the metaphase I arrest. Biorientation, however, depends on an interaction between RZZ, Spindly, and possibly other motor activities, that regulate chromosome movement and attachment status.

## Results

### Analysis of the SAC protein localization in oocytes

The localization of SAC proteins was examined in *Drosophila* oocytes. Mature *Drosophila* oocytes, known as stage 14, arrest at metaphase I with the chromosomes clustered together in a single mass, or karyosome, and the centromeres of homologous chromosomes oriented towards opposite poles. Several SAC proteins were detected at kinetochores in stage 14 oocytes, including BUBR1, BUB3, MPS1 and ROD (Figure 1A). In addition to its kinetochore localization, ROD localized to the spindle region between the pole and the centromeres (Figure 1). This redistribution of ROD from the kinetochores to the spindle is termed “streaming” and has been previously observed in mitotic cells [25, 27, 28]. Another RZZ subunit, ZW10 tagged with HA, overlapped with ROD^GFP^ staining, validating that ROD^GFP^ provides an accurate representation of the RZZ complex’s localization (Figure S 1A).

**Figure 1.**
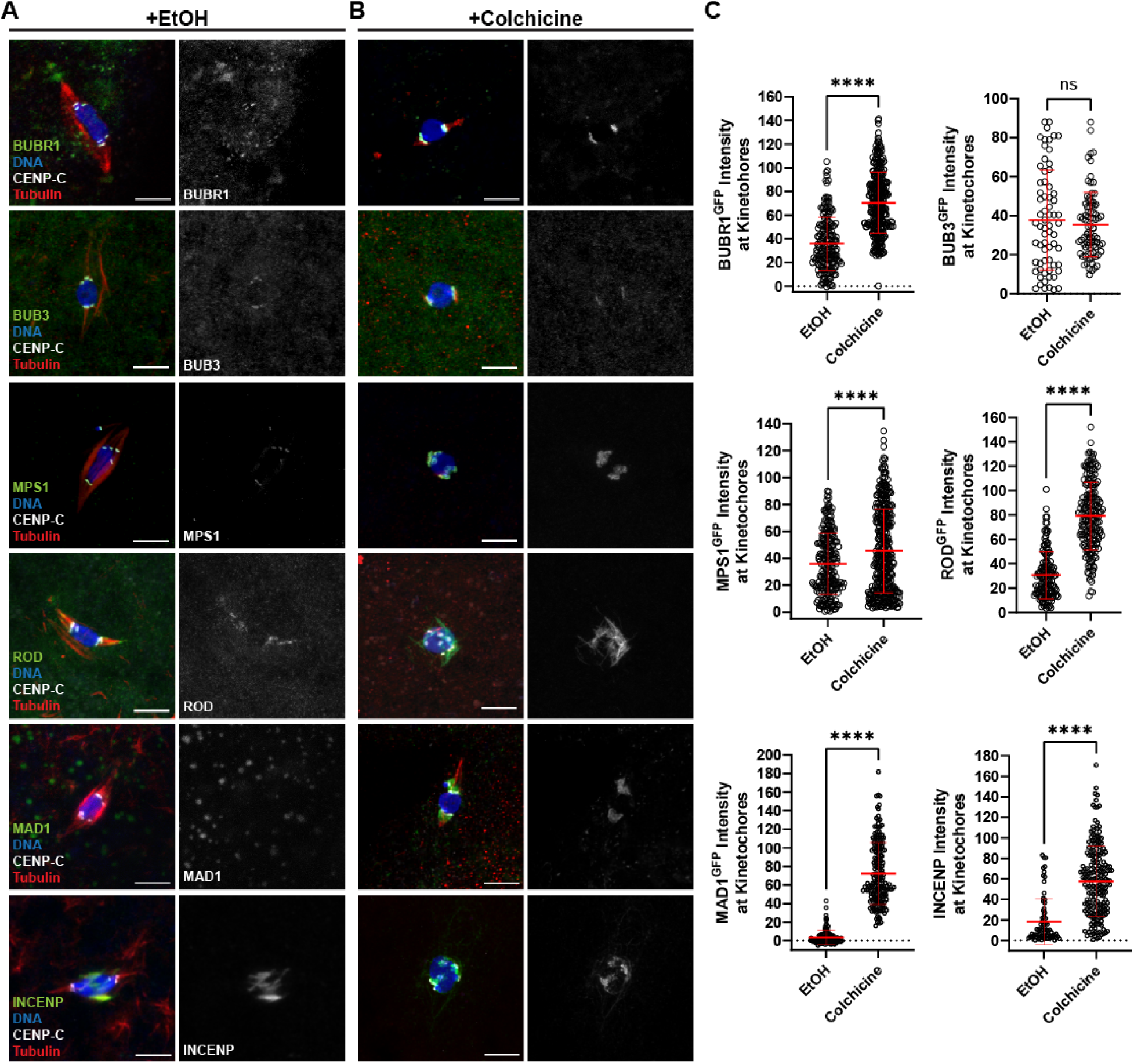
Localization of SAC proteins in oocytes. (A, B) Oocytes were incubated for one hour in Robb’s buffer with 0.5% EtOH (A) or 250 µM colchicine (B). BUBR1, BUB3, MPS1, ROD, and MAD1 were detected using GFP-tagged transgenes, while INCENP was detected using an anti-INCENP antibody (all proteins shown in green), DNA is in blue, CENP-C in white, and tubulin in red. Single channel images show SAC protein localization. All images are maximum intensity projections of z stacks. Scale bars represent 5 µm. (C) Quantification of SAC protein intensity at kinetochores, normalized to background signal (n for EtOH and colchicine = 313 and 174 for BUBR1, 67 and 83 for BUB3, 219 and 356 for MPS1, 139 and 184 for ROD, 138 and 185 for MAD1, and 74 and 200 for INCENP). Error bars show mean ± s.d.; ****P<0.0001, ns=no significance (unpaired two-tailed t test).

In contrast to the above proteins, we did not detect MAD1 in stage 14 oocytes (Figure 1A). Therefore, we attempted to increase SAC signaling by incubating stage 14 oocytes in colchicine for 60 minutes to remove nearly all microtubules. Following this treatment, robust kinetochore localization of MAD1 was observed in all oocytes (Figure 1B). Because colchicine induced a change in MAD1 localization, we tested if the localization of other checkpoint protein was increased by loss of microtubules. BUBR1 and MPS1 intensity increased in colchicine, suggesting their levels are sensitive to the SAC (Figure 1C). We did not, however, observe a SAC response with BUB3 localization.

A more dramatic response was observed with ROD. In colchicine-treated oocytes, ROD expanded around the kinetochores (Figure 1B). In some cases, ROD expanded to surround the karyosome and individual kinetochore foci could no longer be observed. This behavior of the RZZ complex likely corresponds to expansion of the fibrous corona, observed in miotic cells that lack kinetochore-microtubule (KT-MT) attachments [25, 29]. This expansion is thought to facilitate microtubule capture by increasing the surface area of the kinetochore which can contact the microtubules. Thus, in *Drosophila* meiotic oocytes, the fibrous corona expands when microtubules are absent.

Activation of the SAC is also associated with elevated Aurora B activity [30]. In *Drosophila* oocytes, the chromosomal passenger complex (CPC) is recruited to the chromosomes but then moves to the central spindle in prometaphase I oocytes [31]. CPC proteins are not detected at the meiotic centromeres in fixed samples [31, 32]. In colchicine-treated oocytes, however, the CPC component INCENP localized to the chromosomes and the centromeres (Figure 1B, C). These results suggest that, like MAD1, colchicine treatment causes increased localization of the CPC to the centromeres.

### Effect of kinetochore-microtubule attachments on the SAC response

In all colchicine-treated oocytes, MAD1 localization was consistent and observed on all kinetochores, suggesting a global SAC response to the absence of microtubules. To investigate the mechanism of SAC activation, we examined MAD1 localization in oocytes where end-on attachments or tension was disrupted. NDC80 is required for end-on attachments, so nearly all attachments in *Ndc80^RNAi^*oocytes are lateral [22]. MAD1 was not detected in *Ndc80^RNAi^* oocytes unless they were treated with colchicine, suggesting that the SAC is insensitive to the loss of end-on attachments (Figure 2A-C). We also examined MAD1 in a *mei-218* mutant (*mei-218^null^*), which has a high error rate and lacks tension on the kinetochores due to the absence of chiasmata [33]. MAD1 was not detected in *mei-218^null^*oocytes unless they were treated with colchicine, suggesting that loss of tenson does not induce a SAC response (Figure 2A, B, D). The absence of MAD1 in the untreated *mei-218^null^* oocytes is consistent with the prior observation that oocytes lacking chiasma bypass the metaphase I arrest and precociously enter anaphase in stage 14 oocytes [33]. These results show that the SAC is only activated in the absence of microtubules and does not respond to the lack of stable attachments, biorientation errors, or tension.

**Figure 2.**
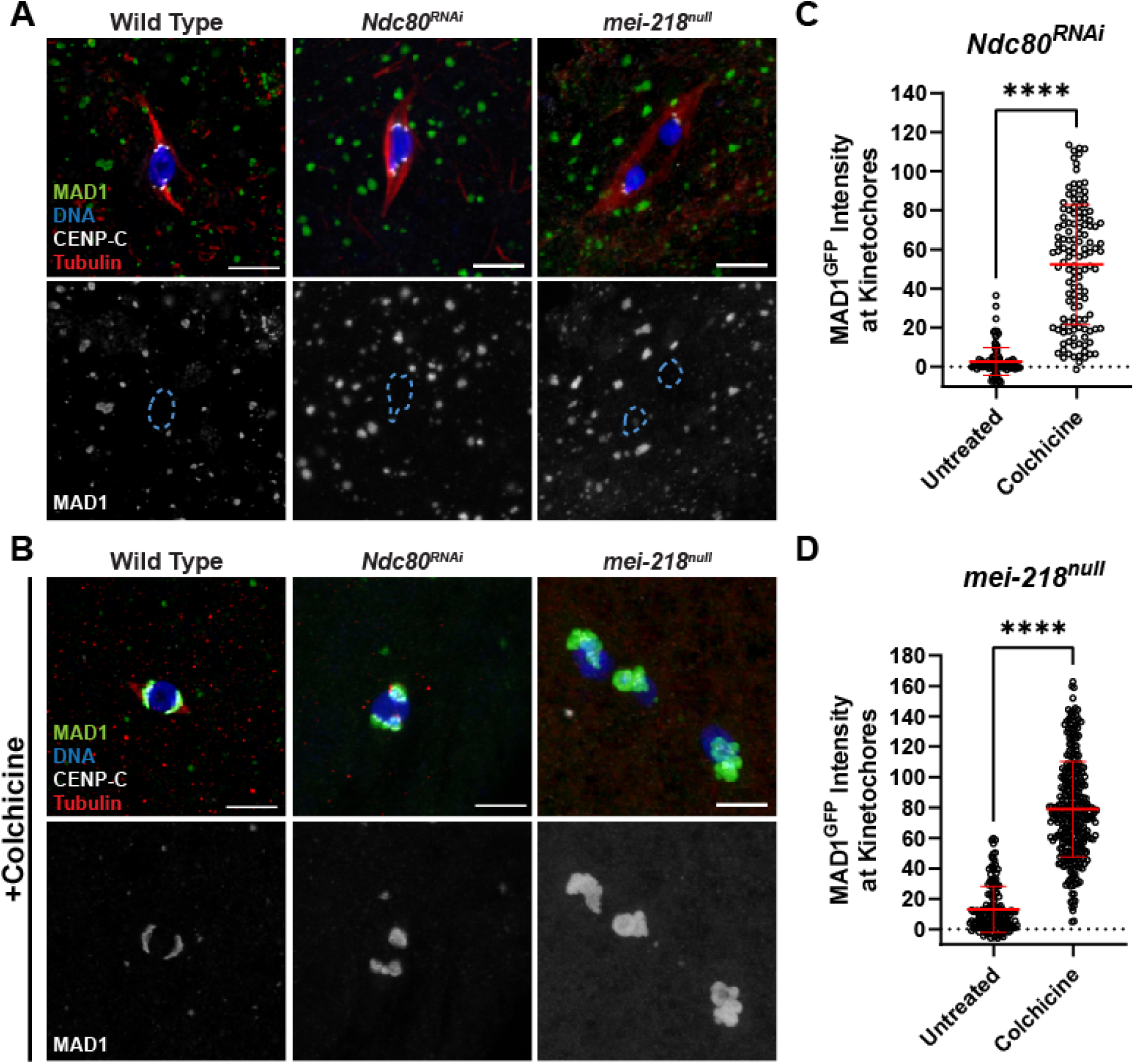
Induction of MAD1 localization is independent of tension and attachment status. (A, B) MAD1^GFP^ localization in wild-type, *Ndc80^RNAi^*, or *mei-218^null^* oocytes that were either untreated (A) or treated with 250 µM colchicine (B). MAD1^GFP^ is shown in green, DNA in blue, CENP-C in white, and tubulin in red. Single channel images (bottom) show MAD1^GFP^ with karyosome outlines in dashed blue in (A). All images are maximum intensity projections of z stacks. Scale bars represent 5 µm. (C, D) Quantification of MAD1^GFP^ intensity at kinetochores, normalized to background GFP signal (n =106 and 127 kinetochores for (C) and 158 and 311 kinetochores for (D)). Error bars show mean ± s.d.; ****P<0.0001 (unpaired two-tailed t test).

### Localization of checkpoint proteins in Drosophila oocytes depends on SPC105R

The outer kinetochore protein KNL1/SPC105R is involved in the recruitment of several checkpoint proteins in mitotic cells [34, 35]. Therefore, we investigated the role of SPC105R, the *Drosophila* homolog of KNL1, in the recruitment of MAD1, ROD, MPS1, and the CPC to kinetochores in *Drosophila* oocytes. To test if SPC105R is required for the recruitment of INCENP and MAD1, we treated *Spc105R^RNAi^* oocytes with colchicine. Colchicine-induced kinetochore localization of both INCENP and MAD1 depended on SPC105R (Figure S 2A-D). However, both still localized to the chromosomes. In the absence of SPC105R, MAD1 moved to the pericentric regions adjacent to the centromeres, while INCENP localized to all chromatin and occasionally to regions adjacent to the chromatin. The observation that MAD1 localized in colchicine-treated oocytes depleted of SPC105R suggests that the oocyte can respond to the loss of microtubules even in the absence of the outer kinetochore.

We also examined localization of INCENP and MAD1 in *Spc105R* deletion mutants to investigate the domains of SPC105R required for the SAC response (Figure S 3). Because *Spc105R* is an essential gene, we used a combination of *Spc105R^RNAi^* and RNAi-resistant transgenes to examine mutants in oocytes. The control for each mutant was an RNAi-resistant wild-type transgene, *Spc105R^B^* (Figure S 3). MAD1 kinetochore localization was reduced in mutants such as *Spc105R^C^*or *Spc105R*^Δ^*^M^* that deleted motifs (MELT, KI, and ExxEED) in the central region of SPC105R (Figure S 2A, B). INCENP kinetochore localization was also lost from the kinetochore in the majority of *Spc105R^C^*or *Spc105R*^Δ^*^M^* oocytes, but due to the diffuse chromatin localization, this was not reflected in significant differences in intensity (Figure S 2C, D). These results suggest that the MELT-like, KI-like, and ExxEED repeats are involved in recruiting MAD1 and the CPC to the kinetochore as part of the SAC response.

MPS1 localization depends on SPC105R, as shown by its drastic reduction in *Spc105R^RNAi^* oocytes (Figure S 4A, B). *Ndc80^RNAi^*oocytes also had a decrease in MPS1 localization, but this was less severe than *Spc105R^RNAi^* oocytes, indicating that the majority of MPS1 is recruited by SPC105R (Figure S 4A, C). Similarly, the *Spc105R^C^*, *Spc105R^ΔM^,* and *Spc105R^ΔExxEED^*mutants had significant decreases in MPS1 localization, but not to the same degree as the *Spc105R^RNAi^* oocytes (Figure S 4B, D).

Each of these mutants retains the C-terminal domain of SPC105R that recruits NDC80 [22]. Thus, these mutants may recruit more MPS1 than *Spc105R^RNAi^* oocytes because NDC80 recruits some MPS1. Therefore, these results suggest that MPS1 is recruited by domains within both SPC105R and NDC80.

ROD localization also depends on SPC105R, as shown by its reduction in *Spc105R^RNAi^* oocytes (Figure 3A, B). In comparison to the other SAC proteins, recruitment of RZZ depends on a smaller and specific domain of SPC105R (Figure S 3, Figure 3, D). Oocytes with a deletion of everything except the KT binding domain (*Spc105R^C^*) or a deletion of all the sequences between the N-terminal and C-terminal domains (MELT-like, KI-like, and ExxEED, termed *Spc105R^ΔM^*) had almost no ROD^GFP^ localization, similar to *Spc105R^RNAi^*oocytes. Furthermore, a deletion of the MELT-KI (*Spc105R^ΔMELT-KI^*), but not the ExxEED domain, had reduced ROD localization. These results suggest that ROD is recruited by the MELT-KI region. Smaller deletions of only the KI and MELT domains (*Spc105R^ΔKI^*and *Spc105R^ΔMELT^*) resulted in significantly reduced ROD^GFP^ localization at the kinetochores, although these intensities were higher than in the *Spc105R^ΔMELT-KI^* deletion. This suggests that ROD is recruited by both the MELT and KI regions.

**Figure 3.**
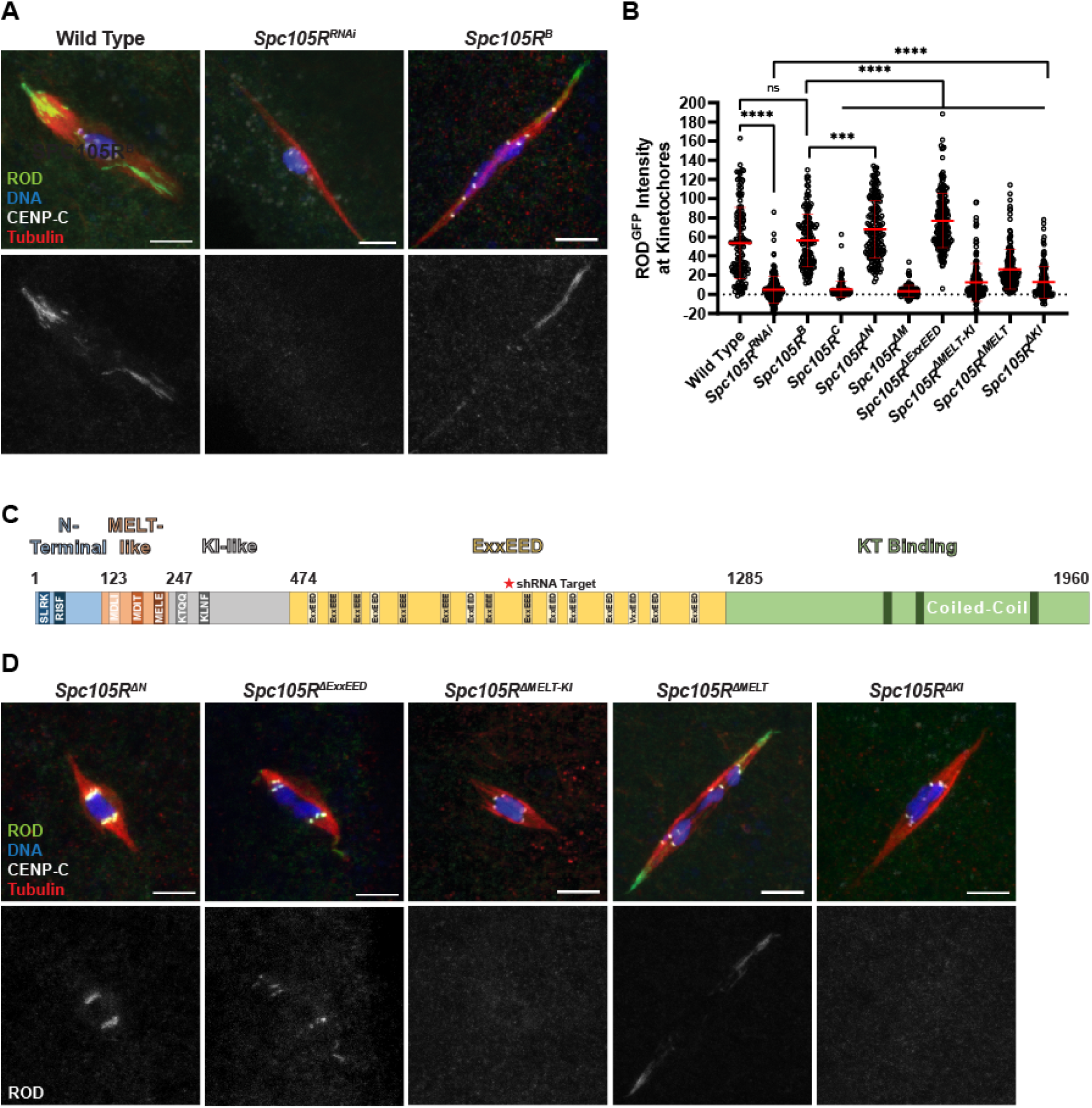
ROD localization depends on the MELT-KI domain of SPC105R. (A) ROD^GFP^ localization in control and an *Spc105R^RNAi^*oocytes, with ROD^GFP^ in green, DNA in blue, CENP-C in white, and tubulin in red. Single channel images (bottom) show ROD^GFP^. (B) Quantification of ROD^GFP^ intensity at kinetochores normalized to background GFP signal in indicated oocytes (from left to right, n = 142, 169, 144, 144, 167, 138, 206, 159, 173, and 146 kinetochores). Error bars show mean ± s.d.; ****P<0.0001, ***P=0.0006, ns=no significance (unpaired two-tailed t test). All *Spc105R* mutants are in an *Spc105R^RNAi^* background targeting the endogenous *Spc105R*. (C) Schematic of SPC105R^B^ domains in *Drosophila melanogaster*. (D) ROD^GFP^ localization in the indicated *Spc105R* mutant oocytes, with ROD^GFP^ in green, DNA in blue, CENP-C in white, and tubulin in red. Single channel images (bottom) show ROD^GFP^. All images are maximum intensity projections of z stacks. Scale bars represent 5 µm.

We also tested if the localization of ROD and MPS1 depended on other SAC proteins. ROD (Figure 4A, B) and MPS1 (Figure 4C, D) localization was significantly reduced in both *Bub3^RNAi^* and *BubR1^RNAi^* oocytes. The more significant reductions that were observed with *Bub3* knockdown could be due to less efficient *BubR1^RNAi^*(see Methods), or redundancy between BUBR1 and BUB1. We also tested the effect of *Cyclin A* knockdown because it may regulate MPS1 localization [36]. *CycA^RNAi^*oocytes had a significant reduction in MPS1 localization, although not as strong as *Bub3^RNAi^*, and a less significant reduction in ROD localization. Furthermore, MPS1 and ROD localization was inter-dependent, although there was a milder reduction of ROD in *Mps1^RNAi^*. These results suggest ROD and MPS1 localization depends on both SAC proteins like BUB3 and Cyclin A.

**Figure 4.**
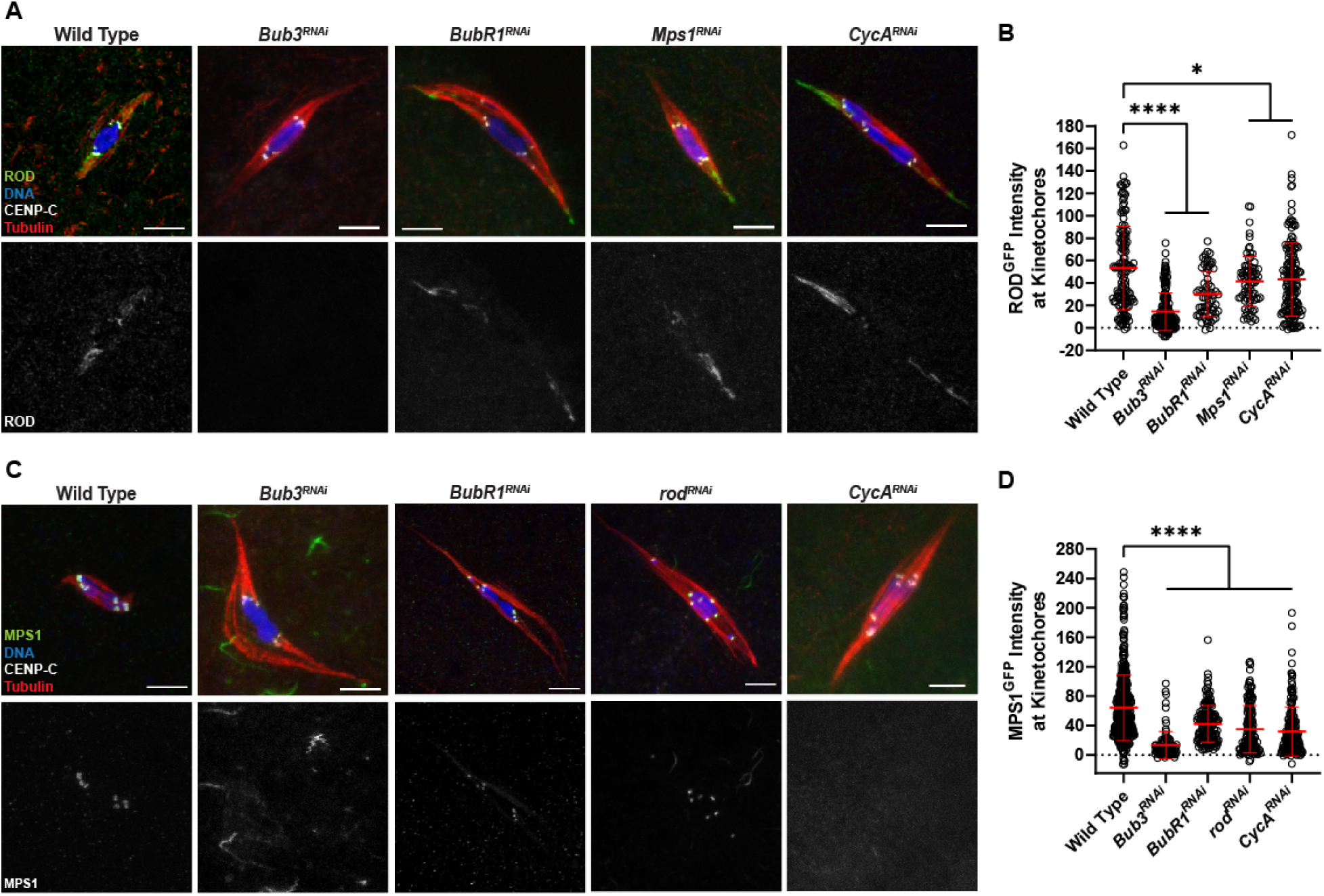
ROD and MPS1 localization depend on SAC genes. (A) ROD^GFP^ localization in wild-type, *Bub3^RNAi^*, *BubR1^RNAi^*, *Mps1^RNAi^*, and *CycA^RNAi^* oocytes with ROD^GFP^ in green, DNA in blue, CENP-C in white, and tubulin in red. Single channel images (bottom) show ROD^GFP^. (B) Quantification of ROD^GFP^ intensity at kinetochores, normalized to background GFP signal in indicated oocytes (from left to right, n = 142, 170, 66, 76, and 155 kinetochores). Error bars show mean ± s.d.; ****P<0.0001, *P≤0.0123 (unpaired two-tailed t test). (C) MPS1^GFP^ localization in wild-type, *Bub3^RNAi^*, *BubR1^RNAi^*, *rod^RNAi^*, and *CycA^RNAi^* oocytes with MPS1^GFP^ in green, DNA in blue, CENP-C in white, and tubulin in red. Single channel images (bottom) show MPS1^GFP^. (D) Quantification of MPS1^GFP^ intensity at kinetochores, normalized to background GFP signal in indicated oocytes (from left to right, n = 455, 97, 149, 180, and 152 kinetochores). Error bars show mean ± s.d.; ****P<0.0001 (unpaired two-tailed t test). All images are maximum intensity projections of z stacks. Scale bars represent 5 µm.

### RZZ and MPS1, but not other SAC proteins, are required for meiosis I biorientation

To determine which SAC genes are required for biorientation during meiosis I, we used fluorescent in-situ hybridization (FISH) probes against the pericentromeric regions of the 2^nd^, 3^rd^, and X chromosomes. Biorientation was defined as two homologous centromeres at opposite ends of the karyosome (Figure 5A). A mono-orientation defect was defined as the two homologous centromeres being in the same half of the karyosome. Oocytes from females expressing *rod^RNAi^* or *Mps1^RNAi^* had increased chromosome mono-orientation (Figure 5B). The results with *Mps1* are consistent with previous studies showing nondisjunction using fertile allele combinations of *Mps1* [37, 38]. We also tested *CycA^RNAi^* oocytes, and consistent with a previous report [39], found an elevated level of biorientation defects (Figure 5B).

**Figure 5.**
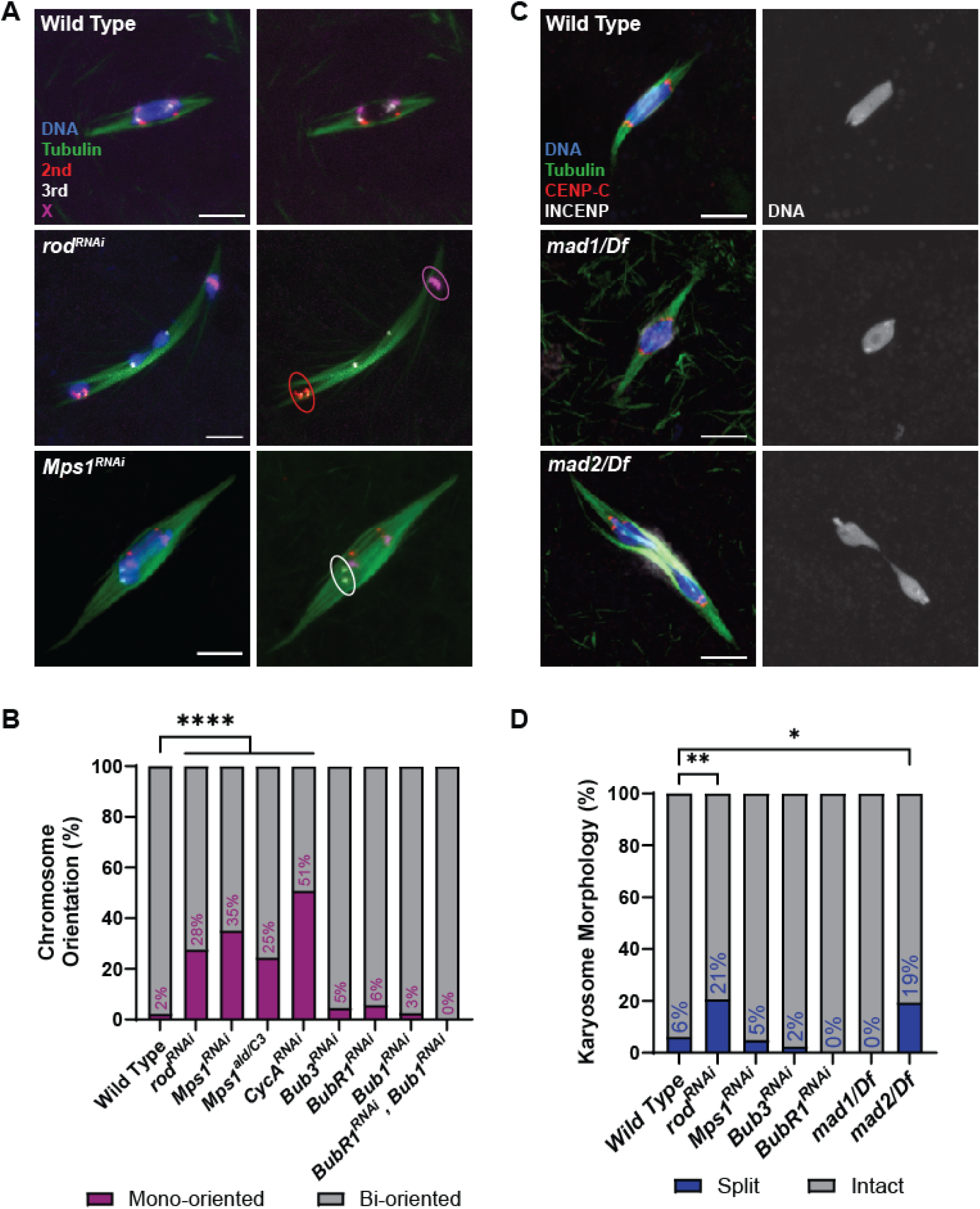
Meiotic chromosome alignment in SAC RNAi oocytes. (A) Representative images depicting chromosome orientation in wild-type, *rod^RNAi^*, and *Mps1^RNAi^*oocytes. Images show DNA (blue), tubulin (green), the X chromosome (magenta), the second chromosome (red), and the third chromosome (white). Tubulin and FISH probes are also shown in a separate channel, with examples of mono-oriented chromosomes circled. All images are maximum intensity projections of z stacks. Scale bars represent 5 μm. (B) Percent of chromosomes per oocyte with the indicated orientation (from left to right, n = 132, 65, 114, 94, 65, 66, 87, 39, and 30 pairs of chromosomes). ****P<0.0001 (Fisher’s exact test). (C) Examples of wild-type, *mad1^1^,* and *mad2^EY21687^* mutant oocytes, with tubulin in green, DNA in blue, CENP-C in red, and INCENP in white. Single channel images show DNA. (D) Percent of oocytes with the specified karyosome morphology (from left to right, n=114, 102, 103, 41, 31, 32, and 31 oocytes). **P=0.0020, *P=0.0375 (Fisher’s exact test).

In contrast, *BubR1^RNAi^*, *Bub1^RNAi^, and Bub3^RNAi^*oocytes had insignificant frequencies of mono-orientation (Figure 5B). Similarly, we have previously shown that flies with knock down of *Bub1* by RNAi are fertile and do not have increased nondisjunction [31]. BUBR1 is required for sister chromatid cohesion protection in *Drosophila*, however, this may have not been detected by our FISH assay at metaphase I [22, 40]. Mutants of *mad1* and *mad2* are viable and fertile [41, 42], so we used genetic crosses to measure the frequency of nondisjunction (see Methods). Neither mutant had a significant increase in nondisjunction (Table 1). We also used a balancer chromosome to suppress crossing over on the X chromosome (*Bwinscy*), thus testing the role of *mad1* and *mad2* in an achiasmate system [43]. Only a mild increase in nondisjunction was observed. These results suggest that most of the SAC proteins are not required for biorientation of homologous chromosomes at metaphase I, while the RZZ complex and MPS1 may be involved in error correction in addition to their roles in the meiotic SAC.

**Table 1:**
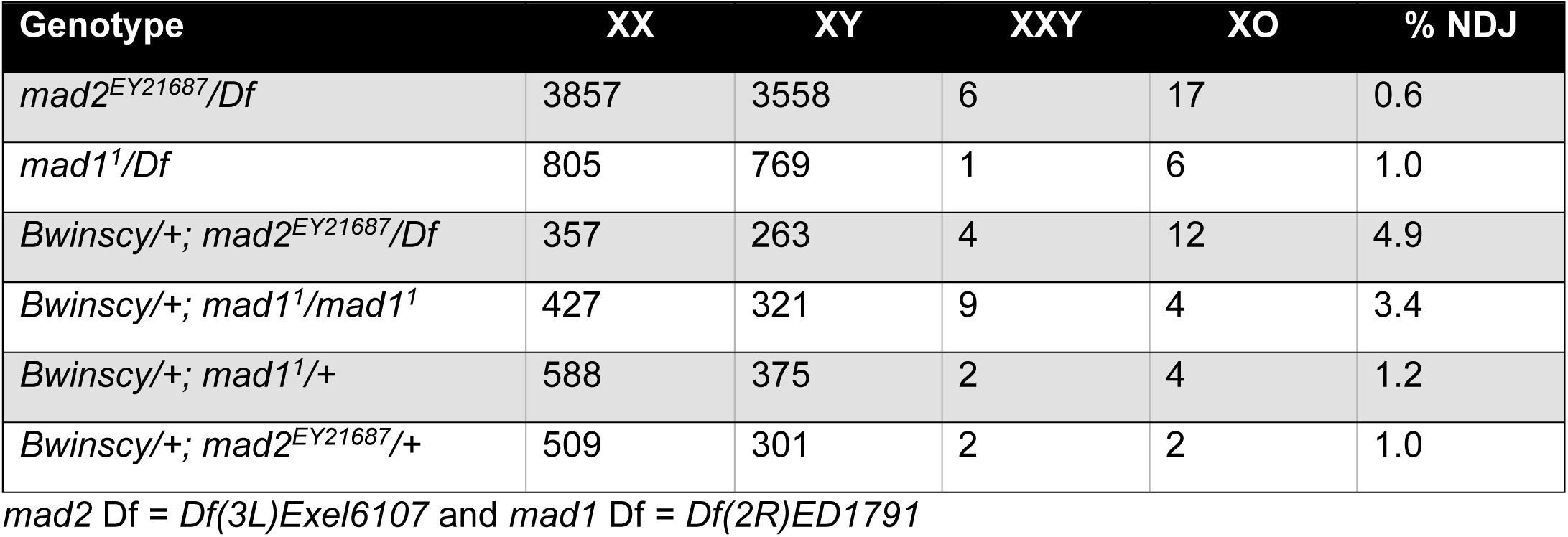
Nondisjunction in *mad* mutants.

Stage 14 oocytes arrest in metaphase I [33, 44]. If the metaphase I arrest depends on the SAC, we would predict that SAC mutants would precociously bypass this arrest and have increased oocytes with split karyosomes. However, this was not observed in most oocytes, suggesting the metaphase I arrest does not depend on SAC activity (Figure 5C, D). The most striking exceptions were *rod^RNAi^* and the *mad2* mutant. The split karyosome phenotype is less than observed in mutants lacking crossing over [22, 33], suggesting these genes could have a minor role in the metaphase I arrest, or in the case of Rod, the precocious anaphase could result from the bi-orientation defects (see below).

### ROD streaming is independent of end-on attachments

MSP1 and ROD may have two distinct functions, one in the SAC and another for biorientation. A striking feature of RZZ is its streaming off the kinetochores, but how streaming contributes to kinetochore attachments and biorientation in meiosis is not known. A small fraction of wild-type oocytes have no ROD streaming, which are likely oocytes in early prometaphase [45]. If streaming is in response to proper attachment of kinetochores to microtubules, we predicted that RZZ would not stream if errors were present, or if end-on attachments could not form. To determine if biorientation errors blocked streaming, we examined ROD in *mei-218^null^* oocytes, in which 90% of crossovers are absent, resulting in frequent chromosome segregation errors [46]. In the absence of crossovers, we still observed precocious anaphase and ROD streaming (Figure 6A,B). Thus, streaming occurs even in the absence of tension and in the presence of attachment errors.

**Figure 6.**
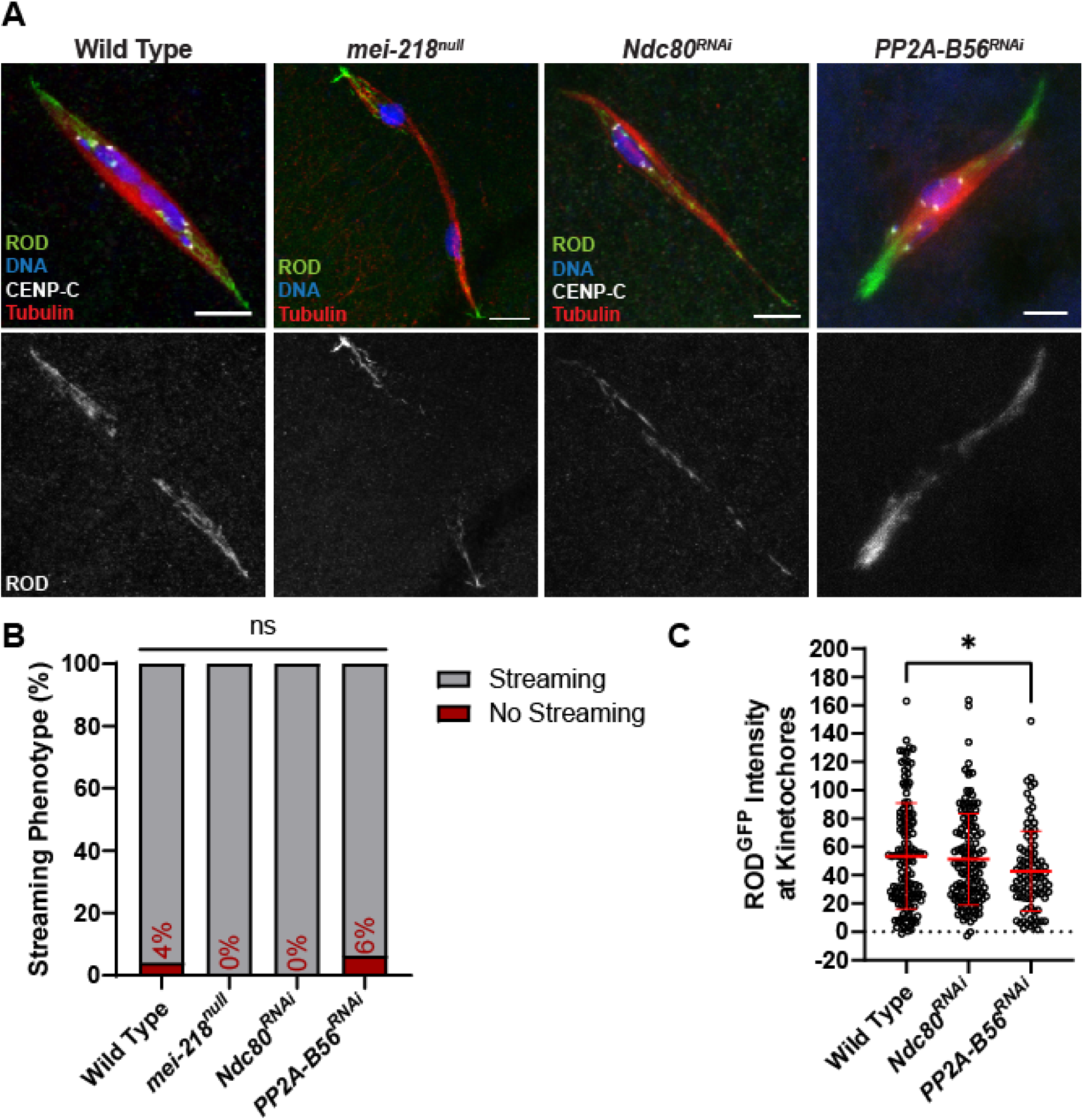
ROD streams in the absence of tension and end-on attachments. (A) ROD^GFP^ localization in wild-type, *mei-218^null^*, *Ndc80^RNAi^*, and *PP2A-B56^RNAi^* oocytes with ROD^GFP^ in green, DNA in blue, CENP-C in white, and tubulin in red. Single channel images (bottom) show ROD^GFP^. All images are maximum intensity projections of z stacks. Scale bars represent 5 µm. (B) Percent of oocytes with ROD^GFP^ streaming in indicated genotypes (from left to right, n = 25, 15, 22, and 16 oocytes), ns=no significance (Fisher’s exact test). (C) Quantification of ROD^GFP^ intensity at kinetochores, normalized to background GFP signal in indicated oocytes (from left to right, n = 142, 139, and 92 kinetochores). Error bars show mean ± s.d.; *P=0.0202 (unpaired two-tailed t test).

To determine if attachment status regulates streaming, we examined ROD in *Ndc80^RNAi^* oocytes, which lack end-on attachments [21]. Despite the lack of end-on attachments, ROD streamed in all *Ndc80^RNAi^* oocytes, and there was no effect on ROD accumulation at the kinetochores (Figure 6A-C). Furthermore, oocytes lacking protein phosphatase 2A-B56 (PP2A-B56), which is also required for the creation of end-on attachments [47], still exhibited ROD streaming (Figure 6A-C). Indeed, there may even be a reduction of ROD at the kinetochore in *PP2A-B56^RNAi^*oocytes. These results demonstrate that end-on attachments are not required for the initiation of RZZ streaming.

### ROD is not required for lateral or end-on attachments

Microtubule nucleation from the kinetochore could depend on RZZ [48]. Thus, the biorientation defects observed in the *rod^RNAi^*oocytes could be caused by the inability to create lateral or end-on attachments, or the creation of inaccurate attachments. If ROD is necessary for making end-on attachments, we would expect that oocytes depleted of *rod* alone would have lateral attachments. However, most attachments in *rod^RNAi^* oocytes were end-on (Figure 7A, B). Interestingly, there was an increase of lateral attachments in *rod^RNAi^* compared to wild-type oocytes. This could be explained if *rod^RNAi^* oocytes are delayed in making end-on attachments, possibly due to frequent biorientation errors. To test whether ROD was necessary for forming lateral attachments, we knocked down both *rod* and *Ndc80* (*rod^RNAi^, Ndc80^RNAi^*). Similar to *Ndc80^RNAi^*, we observed that *rod^RNAi^, Ndc80^RNAi^* oocytes had primarily lateral attachments, suggesting that ROD is not required for making lateral attachments (Figure 7A, B).

**Figure 7.**
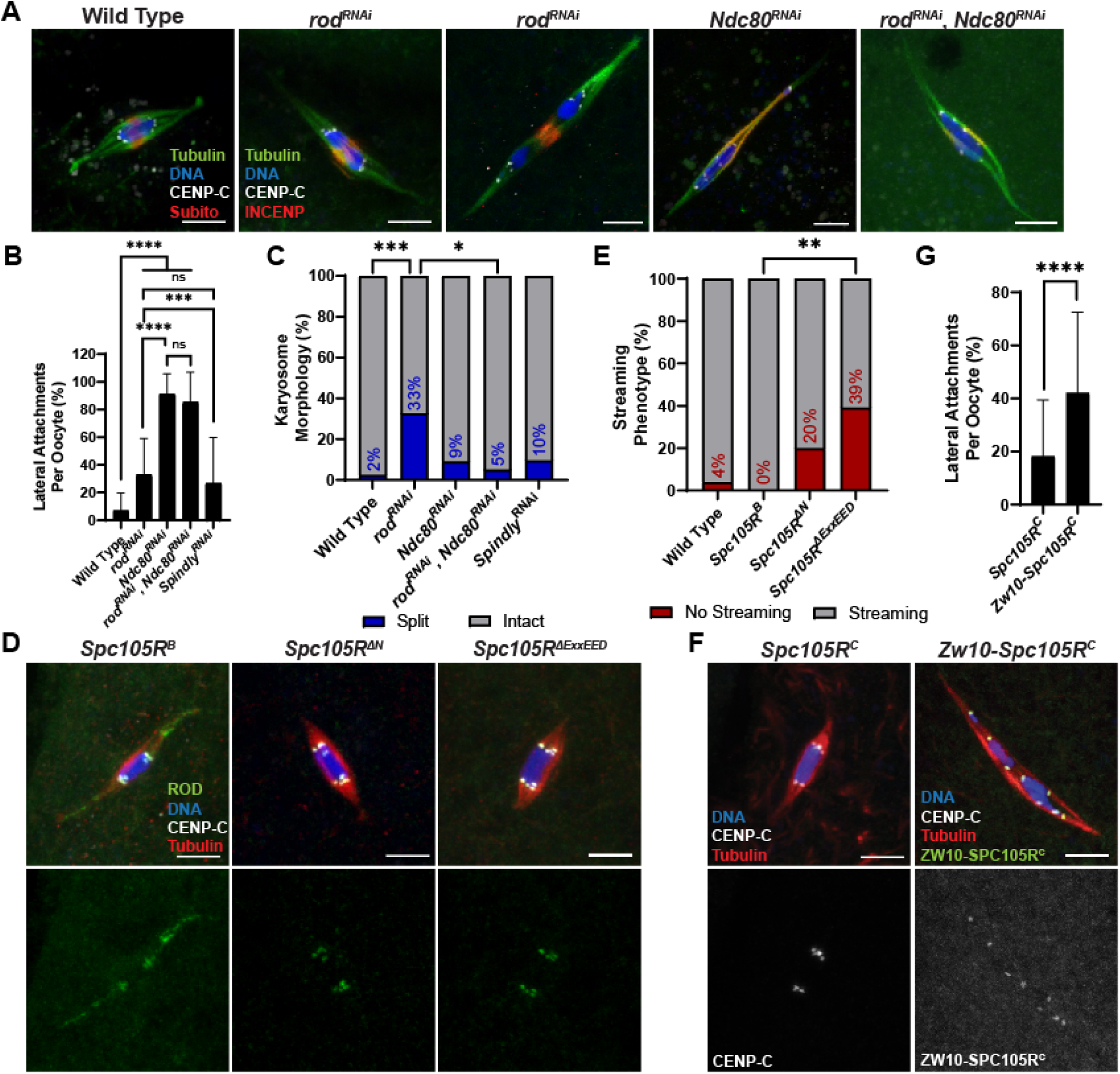
ROD is not required for microtubule attachments. (A) Wild-type; *rod^RNAi^*; *Ndc80^RNAi^*; and *rod^RNAi^*, *Ndc80^RNAi^* oocytes with tubulin in green, DNA in blue, CENP-C in white, and INCENP in red. *Rod^RNAi^*oocyte on the left has an intact karyosome and on the right a split karyosome. (B) Percent of kinetochores per oocyte with lateral kinetochore-microtubule attachments in indicated genotypes (from left to right, n = 40, 46, 30, 20, and 31 oocytes). Error bars show s.d.; ****P<0.0001, ***P=0.0007, ns=no significance (unpaired two-tailed t test). (C) Percent of oocytes with the specified karyosome morphology in the indicated genotypes (from left to right, n = 41, 46, 22, 20, and 31 oocytes). ***P=0.0002, *P=0.0260 (Fisher’s exact test). (D) ROD^GFP^ localization in *Spc105R^B^*, *Spc105R^ΔN^*, and *Spc105R^ΔExxEED^* oocytes with ROD^GFP^ in green, DNA in blue, CENP-C in white, and tubulin in red. Single channel images (bottom) show ROD^GFP^. All *Spc105R* mutants are in an *Spc105R^RNAi^* background targeting the endogenous *Spc105R*. (E) Percent of oocytes with ROD^GFP^ streaming in indicated genotypes (from left to right, n = 25, 20, 20, and 28 oocytes); **P=0.0012 (Fisher’s exact test). (F) *Spc105R^C^* and *Zw10-Spc105R^C^* oocytes (both in an *Spc105R^RNAi^* background) with DNA in blue, CENP-C in white, tubulin in red, and ZW10-SPC105R^C^ in red (right). Single channel images (bottom) show CENP-C (left) or ZW10-SPC105R^C^ (right). (G) Percent of kinetochores per oocyte with lateral kinetochore-microtubule attachments in indicated genotypes (n = 42 and 51 oocytes). Error bars show s.d.; ****P<0.0001 (unpaired two-tailed t test). All *Spc105R^C^* and *Zw10-Spc105R^C^* oocytes are in an *Spc105R^RNAi^* background targeting the endogenous *Spc105R*. All images are maximum intensity projections of z stacks. Scale bars represent 5 µm.

As noted above, *rod^RNAi^*oocytes have an elevated frequency of separated karyosomes. This phenotype was not observed in *rod^RNAi^, Ndc80^RNAi^* oocytes, suggesting that separated karyosome phenotype is caused by the formation of end-on attachments is required for the (Figure 7A, C).

### ROD localization to the kinetochore may delay, but does not prevent, end-on attachments

Because ROD is not required for making lateral or end-on attachments, we reasoned that it could be required for regulating the transition from lateral to end-on attachments. This would align with a model proposed in mitotic cells in which high levels of RZZ at the kinetochores inhibit NDC80 from prematurely establishing end-on attachments [23, 24]. In this model, RZZ streaming allows for the establishment of end-on attachments. If this was true in meiosis, we would expect oocytes in which ROD was exclusively on kinetochores and did not stream to have lateral attachments. This ROD behavior was observed in two *Spc105R* mutants. A deletion of the ExxEED repeats (*Spc105R^ΔExxEED^*) or the N-terminal region (*Spc105R^ΔN^*) had higher ROD intensities at the kinetochores (Figure 3) and less streaming than in *Spc105R^B^* oocytes (Figure 7D, E). However, oocytes with ROD only on kinetochores still exhibited end-on attachments (Figure 7D). These results suggest that the N-terminal region and ExxEED repeats of SPC105R may regulate streaming, and that RZZ streaming is not required for the creation of end-on attachments in *Drosophila* oocytes.

To further test whether end-on attachments are inhibited when the RZZ complex is localized to the kinetochore, we stably targeted RZZ to the kinetochore. The rationale for this experiment is that the recruitment of the RZZ components are interdependent [49]. We fused *Zw10* to the KT-binding C-terminal domain of *Spc105R* (Z*w10-Spc105R^C^*), along with three copies of the HA epitope at the C-terminal end (Figure S 3). The ZW10-SPC105R^C^ fusion protein was targeted to the kinetochores (Figure 7F). When *Zw10-Spc105R^C^*was combined with an shRNA targeting the endogenous *Spc105R* (*Spc105R^RNAi^*), loss of endogenous SPC105R was confirmed cytologically using an antibody against the first 400 amino acids of SPC105R, which is absent in the fusion construct (Figure S 5A). In addition, ROD localized to the kinetochores and did not stream in Z*w10-Spc105R^C^* oocytes, indicating that the RZZ complex was restricted to the kinetochores (Figure S 5B). ROD^GFP^ did not stream even when there was endogenous SPC105R present at the kinetochores. Thus, the ZW10-SPC105R^C^ fusion protein prevented endogenous RZZ streaming from kinetochores. *Zw10-Spc105R^C^* oocytes exhibited an increase in lateral attachments compared to *Spc105R^C^*oocytes (Figure 7G). However, the majority of attachments were still end-on (∼60%), indicating that RZZ may delay, but does not prevent, end-on attachments. Therefore, in agreement with the two *Spc105R* mutants (Figure 7D), streaming of RZZ is not required to establish end-on attachments in *Drosophila* oocytes. Indeed, end-on attachments occur while RZZ is localized to the kinetochore.

### Spindly regulates ROD streaming

Streaming is proposed to be dependent on the microtubule motor protein Dynein [50]. To further examine the role of RZZ streaming and the interaction with Dynein in meiosis, we investigated two dynein-associated proteins previously implicated in RZZ removal from the *Drosophila* kinetochores: Spindly [51], and NudE [52]. We performed FISH on stage 14 oocytes to determine whether these two genes are required for chromosome biorientation (Figure 8A, B). There was only a slight increase in chromosome mono-orientation in *nudE^RNAi^* oocytes compared to wild-type (9% and 0%, respectively). However, 35% of chromosomes were mono-oriented in *Spindly^RNAi^* oocytes, suggesting that Spindly is necessary for accurate meiotic chromosome segregation.

**Figure 8.**
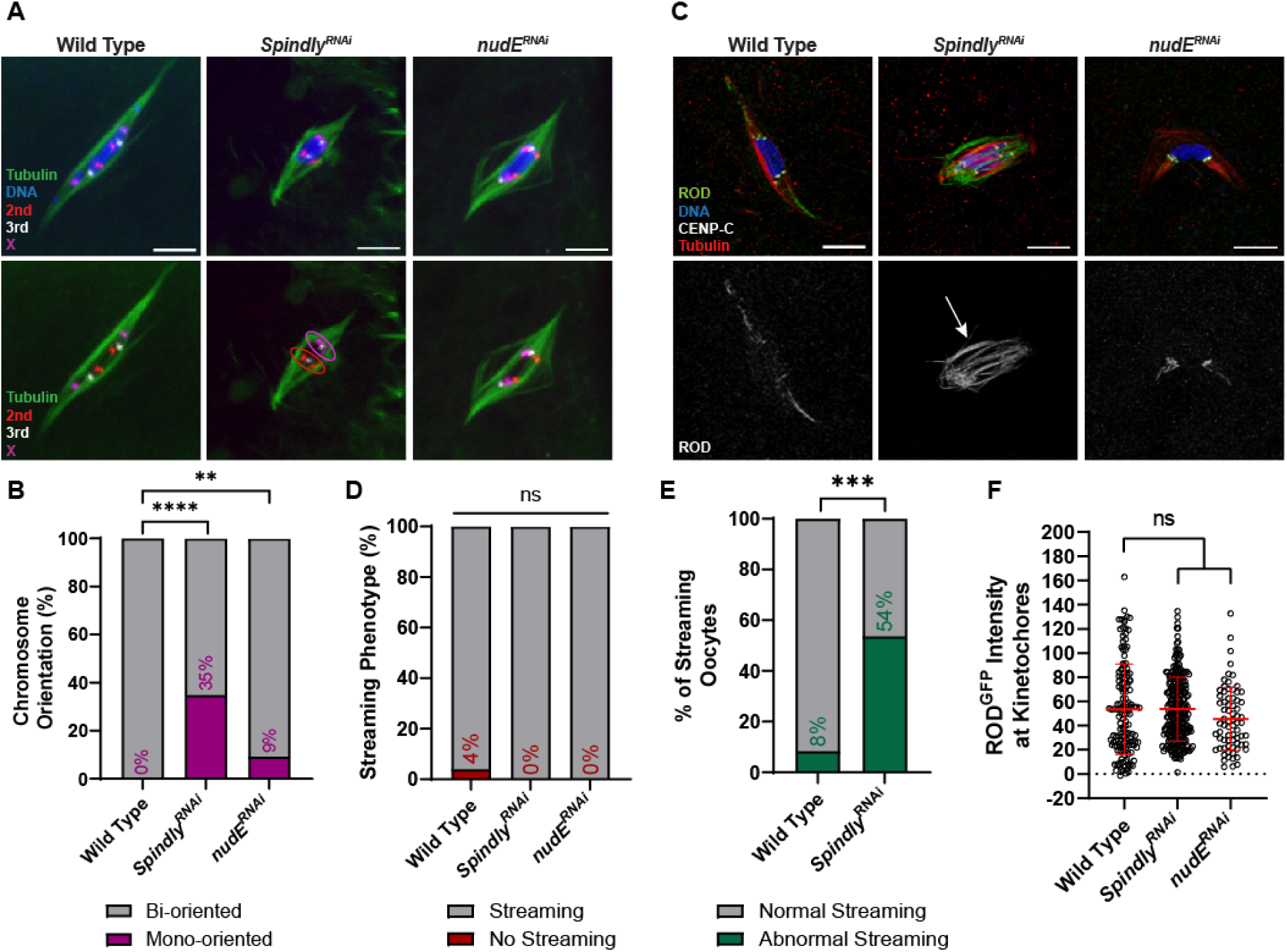
Requirement for Spindly in meiosis and interactions with ROD. (A) Representative images depicting chromosome orientation in wild-type, *Spindly^RNAi^*, and *nudE^RNAi^*oocytes. Images show DNA (blue), tubulin (green), the X chromosome (magenta), the second chromosome (red), and the third chromosome (white). Tubulin and FISH probes are also shown in a separate channel, with examples of mono-oriented chromosomes circled. (B) Percent of chromosomes per oocyte with the indicated orientation (from left to right, n = 86, 66, and 54 pairs of chromosomes); ****P<0.0001, **P=0.0076 (Fisher’s exact test). (C) ROD^GFP^ localization in wild-type, *Spindly^RNAi^*, and *nudE^RNAi^* oocytes with ROD^GFP^ in green, DNA in blue, CENP-C in white, and tubulin in red. Single channel images (bottom) show ROD^GFP^. Arrow points to abnormal streaming in the *Spindly^RNAi^*oocyte. See also Figure S 6A. (D) Percent of oocytes with ROD^GFP^ streaming in indicated genotypes (from left to right, n = 25, 41, and 10 oocytes); ns=no significance (Fisher’s exact test). (E) Percent of oocytes that abnormal streaming, defined as ROD^GFP^ in the central spindle, in wild-type or *Spindly^RNAi^* oocytes (n=24 and 43 oocytes); ***P=0.0002 (Fisher’s exact test). Abnormal streaming is defined as having ROD^GFP^ in the central region between the centromeres.(F) Quantification of ROD^GFP^ intensity at kinetochores, normalized to background GFP signal in indicated oocytes (from left to right, n = 142, 256, and 72 kinetochores. Error bars show mean ± s.d.; ns=no significance (unpaired two-tailed t test). All images are maximum intensity projections of z stacks. Scale bars represent 5 µm.

To determine whether these proteins regulate streaming, we characterized ROD behavior in oocytes depleted of *Spindly* or *nudE* (Figure 8C-E). Both *Spindly^RNAi^*and *nudE^RNAi^* oocytes exhibited ROD streaming (Figure 8D). This was surprising for Spindly, which has been shown to be required for streaming in *Drosophila* S2 cells [51].

However, *Spindly^RNAi^*oocytes also exhibited unusual behavior, notably ROD streaming in the central spindle region between homologous centromeres in approximately 50% of oocytes (Figure 8E, Figure S 6A). Thus, Spindly may be required for proper streaming towards the spindle pole. To confirm that ROD was being efficiently removed and not accumulating at the kinetochores, we measured the intensity of ROD in *Spindly^RNAi^* and *nudE^RNAi^*oocytes (Figure 8F). Both RNAi oocytes had ROD levels similar to those in wild-type oocytes, indicating that ROD can still stream from kinetochores without NudE or Spindly. Therefore, while neither protein is required for RZZ streaming, Spindly may be required for RZZ to stream towards the spindle poles.

RZZ is proposed to recruit Spindly to the kinetochore, which is followed by streaming of both Spindly and RZZ [51, 53]. To examine the localization of Spindly on the meiotic spindle, we used previously described *Spindly-GFP* fusion transgenes [54]. Oocytes expressing wild-type *Spindly* had weak localization throughout the spindle (Figure S 6B). We also examined *Spindly* mutants with deletions of domains implicated in streaming: the N-terminus (a deletion of amino acids 1-228) and the Spindly Box (a deletion of amino acids 229-250). Both mutants had a dominant sterility phenotype when expressed in oocytes using *matα*, suggesting that these mutants interfere with the activity of the wild-type Spindly protein. In contrast to the wild-type oocytes, which had weak localization throughout the spindle, both mutants had pronounced localization near the centromeres. Spindly^ΔN^ localized to the kinetochores and spindle, while Spindly^ΔSB^ primarily localized to the kinetochores (Figure S 6B, C). These results suggest that the Spindly Box domain is important for removal of Spindly from the kinetochores. The robust localization of these mutants was then used to test if ROD is required for Spindly localization, as shown in mitotic cells [51]. Indeed, Spindly was absent in *rod^RNAi^* oocytes, indicating ROD is required for Spindly localization in oocyte meiosis (Figure S 6C, D).

## Discussion

### The function of the SAC in oocytes

We analyzed the localization and meiotic phenotypes of several SAC proteins in *Drosophila* oocytes. Increased localization of MAD1, BUBR1, INCENP, and ROD at the kinetochores following loss of microtubules indicated *Drosophila* oocytes have a functioning SAC. This SAC response was observed even in the absence of a kinetochore. However, a SAC response was not observed following loss of microtubule attachments to the kinetochores, the lack of tension, or the presence of multiple biorientation errors. Thus, the function of the SAC in oocytes may be to promote spindle assembly, rather than proper microtubules attachments to kinetochores [28].

Only two of the SAC proteins we tested were required for biorientation of homologous chromosomes at meiosis I, MPS1 [37, 38] and ROD, suggesting that the SAC is not required for error correction. Similarly, *C. elegans* may correct attachment errors using a mechanism that does not rely on the SAC [55]. The SAC may have a more prominent role in chromosome segregation in mouse oocytes that arrest in meiosis II, where increased aneuploidy has been observed with loss of SAC proteins [56, 57]. However, the evidence comes from a variety of assays that may not be comparable to our FISH experiments, and one study concluded that Mad2 is dispensable for accurate meiotic chromosome segregation except in response to stress [58].

In most cases, the loss of SAC proteins in *Drosophila* oocytes does not result in progression into anaphase (Figure 5) or a rise in APC activity [59]. These observations suggest the SAC may not regulate the APC in metaphase I-arrested oocytes. Instead, the metaphase I arrest in *Drosophila* oocytes may be maintained by a SAC-independent mechanism relying on chiasmata and cohesion, with progression into anaphase I depending on cohesion release. This would explain why *Drosophila* oocytes lacking chiasmata and tension bypass metaphase I arrest and progress into anaphase I [33, 60].

Furthermore, several observations suggest that the SAC is silenced in oocytes, possibly to allow for developmental regulation of the metaphase I arrest, with progression into anaphase occurring only after oocytes enter the oviduct.

Developmentally regulated SAC silencing would explain why ROD streams in wild-type metaphase I-arrested oocytes, even in the absence of end-on attachments or the presence of multiple biorientation errors in *Ndc80^RNAi^*and *PP2A-B56^RNAi^*oocytes [61]. Similarly, MAD2 localizes to centromeres in early (stage 13) but not late (stage 14) stage oocytes [45], and the metaphase I spindle has some anaphase-like characteristics such as high concentrations of the CPC on the central spindle [62].

Yeast meiotic cells silence their SAC to prevent prolonged arrest, indicating this may be a conserved feature of meiotic cells [63].

### Recruitment of checkpoint proteins during meiosis

Our results are consistent with the known role of KNL1/SPC105R in the recruitment of checkpoint proteins to the kinetochore [1]. We previously showed that BUBR1 is recruited by both the MELT-KI and C-terminal regions of SPC105R [22]. MAD1, INCENP, and probably BUB3 depend on the MELT-KI and ExxEED repeat regions of SPC105R, which correspond to the repeated MELT motifs found in vertebrate KNL1. In the absence of SPC105R, recruitment of MAD1 shifts to the peri-centromeric regions. Thus, some components of the SAC do not depend on SPC105R, and there is a SPC105R-independnet SAC response. Human cells may also have two SAC pathways, and one is active in the absence of KNL1 [64]. Thus, centromere components may recruit SAC proteins. In *Drosophila*, this could be facilitated by ZW10, which interacts with and is recruited by centromere protein CAL1 [65].

### Multiple functions and recruitment factors for MPS1 and RZZ

MPS1 was originally discovered with the mutation *ald* as being required for meiotic chromosome segregation [66], and subsequent studies have characterized its role in *Drosophila* meiosis [37, 38] and mouse oocytes [67]. While MPS1 has been shown to be recruited by NDC80 [68–71], two observations show that a significant amount of MPS1 is also recruited by SPC105R. First, we observed a more severe decrease in MPS1 localization upon *Spc105R* depletion than *Ndc80* depletion. Second, we observed a decrease in MPS1 localization in *Spc105R* mutants that still recruit NDC80. These data suggest that MPS1 is recruited by both NDC80 and SPC105R in *Drosophila* oocytes. The SPC105R-dependent MPS1 could function in the SAC while NDC80-dependent MPS1 could regulate attachments [68, 72].

Like MPS1, the RZZ complex has roles in error correction and in the SAC. Like MPS1, our data suggest there are two separate RZZ populations: one that is recruited by BUB3 and BUBR1 and is involved in the SAC and a second that is recruited by the MELT-KI domains of SPC105R and is important for EC. The SAC population of ROD that depends on BUB3 could recruit other checkpoint proteins like MAD1 and MAD2 [73, 74]. These conclusions are based on the observation that unlike *rod^RNAi^*, *Bub3^RNAi^* and *BubR1^RNAi^* oocytes do not have biorientation defects, suggesting that BUB3 and BUBR1 do not recruit the population of RZZ which is involved in error correction. Furthermore, there is a substantial amount of ROD^GFP^ remaining after *Bub3* and *BubR1* are knocked down, arguing that they do not recruit the total population of RZZ at kinetochores. However, the *Spc105R^ΔMELT-KI^*mutant (deleting amino acids 154-840) abolishes nearly all ROD^GFP^ localization and does have biorientation defects [22].

Therefore, the MELT-KI domains of SPC105R are likely involved in recruiting the populations of RZZ involved in EC, independently of BUB3.

The MELT-KI region of SPC105R is highly conserved [75]. Indeed, our results refine observations made in HeLa cells showing that the N-terminal domain of KNL1 recruits RZZ [76, 77]. In *Drosophila* cell lines, a region within the KI-like domain (amino acids 266-384) and BUBR1 are required for RZZ recruitment to kinetochores [78].

Interestingly, this study also showed that the recruitment of RZZ by SPC105R could promote chromosome movement towards the spindle poles. Like this study, we found the KI domain recruits the majority of RZZ. It remains to be determined, however, why both the MELT and KI domains are required to recruit RZZ and for biorientation in meiosis I [22].

ROD and MPS1 localization was reduced in *CycA^RNAi^* oocytes, and *CycA^RNAi^* oocytes have biorientation defects in *Drosophila* oocytes [39]. Indeed, Cyclin A has been shown to regulate KT-MT attachments in mammalian cells [79, 80] and in *Drosophila* in mitotic cells, Cyclin A/CDK1 activates Polo kinase, which has recently been shown to regulate Spindly and prevent it from inhibiting RZZ [24]. One explanation for these results is that Cyclin A is required for the functions of ROD and MPS1 associated with ensuring the accuracy of biorientation.

### The role of RZZ localization in attachment stabilization

Previous studies in *C. elegans, Drosophila,* and human mitotic cells have suggested a two-part model for how RZZ regulates KT-MT attachments [23, 24, 81, 82]. First, RZZ interacts with NDC80 to inhibit end-on attachments. Second, Spindly-dependent RZZ streaming alleviates RZZ inhibition to allow for the formation of end-on attachments. Three of our observations are consistent with this hypothesis. First, knockdown of *Spindly* resulted in a decrease in end-on and an increase in lateral attachments. This would be expected if Spindly negatively regulates RZZ. Second, Z*w10* fused to the C terminus of *Spc105R* led to complete RZZ retention on the kinetochores and increased lateral attachments. Third, we observed an increase in karyosome separation in *rod^RNAi^*oocytes, which could be due to precocious stabilization of end-on attachments [22, 83]. These results suggest that the RZZ complex at the kinetochore has a role in preventing or delaying end-on attachments.

However, some of our results are not consistent with this model. Oocytes expressing *Zw10-Spc105R^C^* primarily had end-on attachments. A similar observation was made with two mutants (*Spc105R^ΔExxEED^*and *Spc105R^ΔN^*) that had reduced ROD streaming but still formed end-on attachments. Therefore, if KT-localized RZZ inhibits end-on attachments, this can be overridden in oocytes. Finally, ROD streaming was observed in the absence of end-on attachments. Thus, physical removal of RZZ (or streaming) may not be required for the formation of end-on microtubule attachments, and alternative mechanisms for regulating this population of RZZ may exist. For example, a conformational change in RZZ may affect its regulation of attachments [23]. Understanding the relationship between the localization of the RZZ complex and its effects on microtubule attachments is crucial for analyzing its role in promoting accurate chromosome segregation during female meiosis.

### The role of Spindly in meiotic RZZ streaming and homolog biorientation

Spindly has a conserved role as a Dynein adaptor [84], and we have shown that RZZ recruits Spindly to kinetochores in *Drosophila* oocytes. Spindly has been shown in multiple organisms to recruit Dynein, resulting in removal of the RZZ complex from the kinetochores to stabilize microtubule attachments [24, 53, 81, 85]. This activity has been used to explain why depletion of RZZ can suppress defects in mitotic cells depleted of Spindly [23, 24, 81]. However, the role of Spindly in meiosis has not been reported.

Spindly is required for biorientation of homologous chromosomes during oocyte meiosis. The failure to remove RZZ from the kinetochores could be the reason for the meiotic biorientation defect. Contrary to the expectations from data in mitotic cells described above, however, knocking down *Spindly* did not abolish ROD streaming.

Instead, we found that ROD displayed abnormal behaviors in oocytes depleted of *Spindly*, such as streaming in the opposite direction and localizing in the central spindle region between kinetochores. Therefore, it is likely that other proteins are involved in the removal of RZZ from the kinetochores in oocytes.

In the absence of Spindly in HeLa cells, alternative mechanisms are activated to remove kinetochore proteins [86]. One candidate is CENP-E, which is a plus-end directed motor required for the fibrous corona [87], and is required for homolog biorientation in *Drosophila* oocytes [21]. Given that RZZ can move SPC105R oligomers [88], and our observation that streaming off the kinetochore may not affect attachment status, an important role of RZZ may be to move kinetochores. Rather than being necessary for the direct removal of RZZ from the kinetochores, Spindly, CENP-E, and RZZ might form a complex capable of bidirectional movement of the kinetochores [89]. Understanding how RZZ regulates biorientation may depend on reconciling the relative importance of two activities, RZZ moving kinetochores with motors like Dynein and CENP-E, and RZZ blocking or delaying end-on attachments.

## Materials and Methods

### Drosophila Genotypes and Expression of Transgenes and shRNAs

The transgenes and mutants used in this study are listed in Table 2. The localization of SAC proteins was examined in *Drosophila* oocytes using GFP fusion transgenes, most under the control of their own promoter. To examine MPS1, we used a construct used extensively in cell culture but modified for making transgenics (*P{EGFP-Mps1.F}* = *Mps1^GFP^*) [90, 91]. To visualize the RZZ complex in oocytes, we used a construct where ROD was tagged with Green Fluorescent Protein (GFP) at the N-terminus and under the control of its own promoter (*P{gEGFP-rod}* = *rod^GFP^*) [25] (Figure S 1A). Expression of either transgene resulted in very low levels (∼0.25%) of X-chromosome nondisjunction (Table S1), which was comparable to wild-type strains [92]. This is consistent with previous studies showing that this transgene can rescue the lethality of *rod* mutants [25, 93].

**Table 2.**
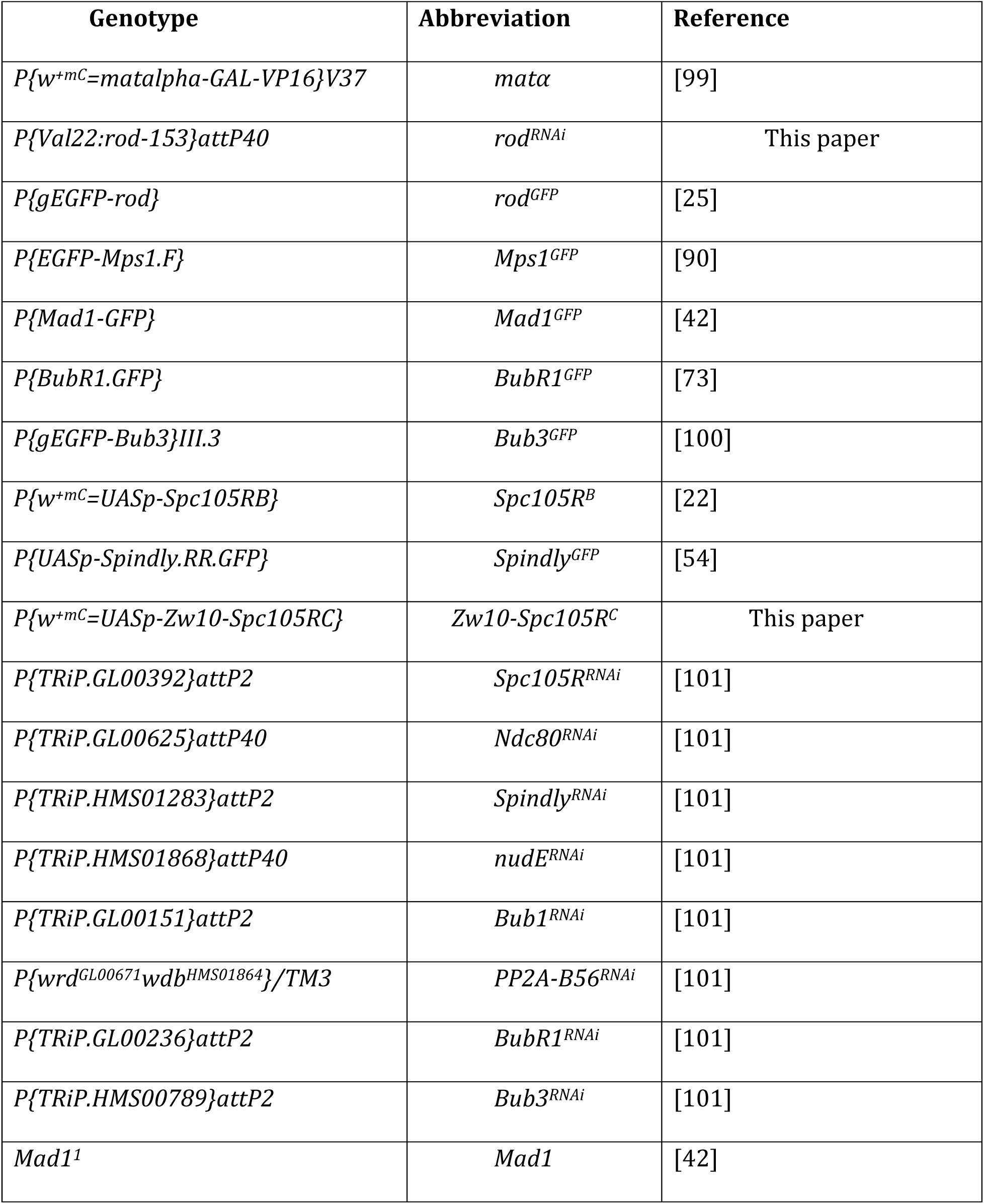
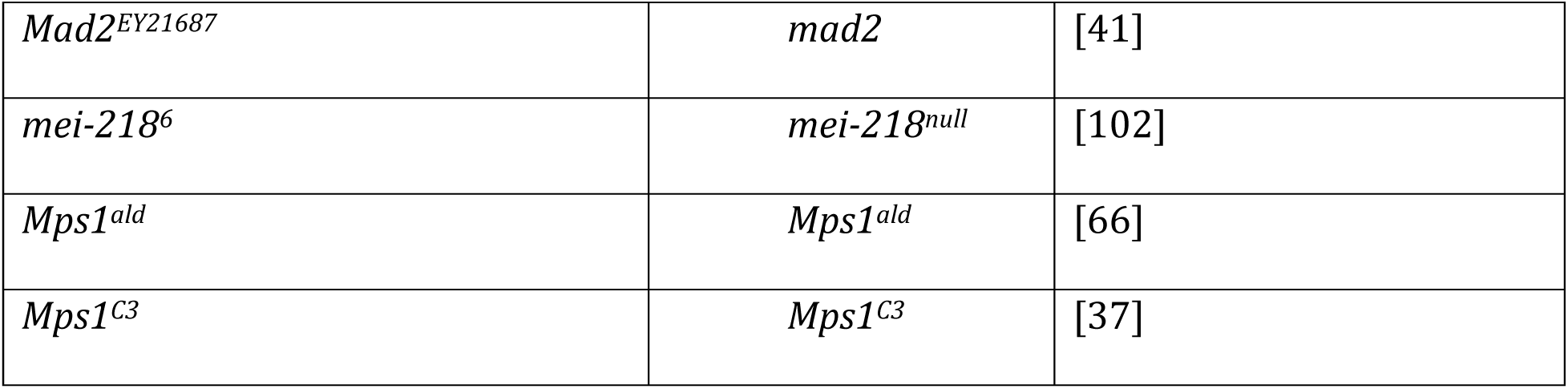
*Drosophila* genotypes used in this work.

The UAS/GAL4 system was utilized to drive expression of many of the transgenes in oocytes. *Matα* was used to induce expression following DNA replication and early pachytene and throughout meiotic prophase. A fusion between the C-terminal domain of *Spc105R* and the entire coding region of Z*w10* was generated by GenScript in pENTR4. The C-terminal fragment of SPC105R has been characterized previously and is sufficient for kinetochore localization [22]. The fusion gene was transferred into pPWH, which fused the coding region to a 3XHA tag at the C-terminal end. This was injected into embryos for germline transformation by Model Systems Injections.

### RNA Extraction and Quantification of RNAi Knockdown

Among three shRNA lines targeting *rod*, one (*P{Val22:rod-153}1attP40,* = *rod^RNAi)^*) was chosen because it had the strongest phenotype and most severe mRNA knockdown (4% of the wild-type). Expression of *rod^RNAi^*using *matα* produced sterile females and ubiquitous expression caused lethality. To confirm the efficacy of the *rod^RNAi^*in oocytes, we examined ROD^GFP^ in oocytes expressing the *rod* shRNA. The intensity of ROD^GFP^ at the kinetochores in *rod^RNAi^*oocytes was reduced to nearly background levels (Figure S 1B, C), indicating that the *rod* shRNA provides a strong knockdown of *rod*. Similarly, the intensity of MPS1^GFP^ at the kinetochores in *Mps1^RNAi^* oocytes was significantly reduced (Figure S 1D, E).

RT-qPCR was performed to determine knockdown of mRNAs for each shRNA. After the parental flies were crossed, mature female offspring with the correct genotype were collected into yeasted vials for three days. These flies were then ground in 1X phosphate-buffered saline (PBS) and filtered through meshes to isolate the stage 14 oocytes. 1 mL of TRIzol reagent was added and RNA was extracted per the manufacturer’s instructions (Thermo Fisher Scientific). Taqman RT-qPCR was performed to quantify RNAi knockdowns. 2 μg of isolated RNA was reverse transcribed into cDNA using the High Capacity cDNA Reverse Transcription Kit (Applied Biosystems). Afterwards, qPCR was performed using a TaqMan Assay (Thermo Scientific) with four replicates per reaction. The efficacy of RNAi knockdowns has been published for *Ndc80 GL00625* = 6% remaining*, Spc105R GL00392* = 13% remaining [21]*, Bub1 GL00151* = 2% remaining [31], *BubR1 GLV21065* = 22% remaining, *BubR1 GL00236*= 27% remaining, and *Bub3 HMS00789* = 0.7% remaining.

### Measuring X-Chromosome Nondisjunction

To measure nondisjunction of the X chromosome, virgin females with the desired genotype were crossed to males with a dominant *Bar* mutation on the Y chromosome (*yw/B^S^Y*). This mutation led to a visible bar-eyed phenotype where the eye was thinner than wild-type eyes. The normal segregation of chromosomes would lead to female offspring (XX) with wild-type eyes and males (XY) with bar eyes. If a nondisjunction event occurred in the X chromosome, the following sex chromosome combinations were possible: XO, XXY, XXX, and YO. XO and XXY were viable and resulted in normal-eyed males and bar-eyed females, respectively. XXX and YO were lethal. To calculate nondisjunction rates while taking into account the lethal genotypes, the following equation was used: 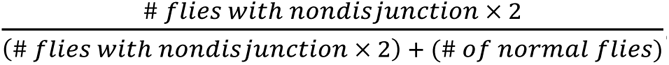. This rate determined whether a particular genotype resulted in increased meiotic nondisjunction.

### Cytology of Stage 14 Oocytes

Cytology of stage 14 oocytes was performed as previously described [94]. After the parental flies were crossed, mature female offspring with the correct genotypes were collected into yeasted vials for one to two days. These flies were then broken open in a blender while in 1x modified Robb’s buffer and filtered through meshes to isolate the stage 14 oocytes. If a colchicine treatment was needed, oocytes were incubated with 150 μM colchicine or 0.5% EtOH in modified Robb’s buffer for one hour to remove all microtubules. The oocytes were fixed with 5% formaldehyde and heptane before being rinsed in 1X PBS. The oocytes were then rolled between the frosted surface of a glass slide and a cover slip to remove the membranes and nutated for two hours in PBS/1% Triton X-100 to make them permeable to antibody staining. Afterwards, oocytes were washed in PBS/0.05% Triton X-100 and blocked for one hour in blocking buffer (0.5% BSA and 0.1% Tween-20 in 1X PBS). Subsequently, primary antibodies were added to the oocytes while the secondary antibodies were added to *Drosophila* embryos to remove any nonspecific binding. After nutating overnight at 4°C, the oocytes were washed in blocking buffer four times at room temperature for 15 minutes. The secondary antibodies were added and the oocytes were nutated for 3-4 hours at room temperature. The oocytes were then washed in blocking buffer with Hoechst33342 (10 μg/mL) to stain the DNA. The oocytes were washed twice more in blocking buffer and then were ready for mounting and imaging.

The primary antibodies used in this paper were tubulin monoclonal antibody DM1A at 1:50 conjugated to FITC (Sigma-Aldrich), tubulin E7 monoclonal antibody at 1:150 [95], rat anti-INCENP at 1:600 [96], guinea pig anti-CENP-C at 1:1000 [97], rat anti-tubulin at 1:300 (Millipore), rabbit anti-Spc105R (1:4000) [98], rabbit anti-GFP (1:200) (Thermo Fisher), rat anti-HA (Roche, 1:50). Multiply preabsorbed secondary antibodies were from Jackson ImmunoResearch, and were conjugated to Alexa 647, Cy3, Alexa 594 or Alexa 488.

To visualize pairs of homologous chromosomes, fluorescent *in situ* hybridization (FISH) was performed. After oocytes fixation, 2X SSC was added and the membranes were removed by rolling the oocytes between glass slides. The oocytes were then nutated in increasing concentrations of formamide solution (20%, 40%, and 50%) before being added to the hybridization solution and probes. The probes were synthesized by IDT, and recognize the 359 bp repeats on the X chromosome (labeled with AlexaFluor 594), the AACAC repeats on the second chromosome (labeled with Cy3), or the dodeca repeats on the third chromosome (labeled with Alexa 647). This was followed by incubation in 80°C for 20 minutes and then 37°C overnight. The next day, the oocytes were washed in 50% formamide solution twice at 37°C and 20% formamide once at room temperature. The oocytes were then rinsed three times in 2X SSCT, once in 2X PBST, and nutated in blocking buffer for 4 hours at room temperature. Afterwards, mouse anti-tubulin-FITC antibody (Sigma) was added (1:50) and the oocytes were nutated overnight at room temperature. The next day, the oocytes were washed in blocking buffer with Hoechst33342 (10 μg/mL). After washing twice more with blocking buffer, the oocytes were mounted and imaged.

### Imaging Oocytes, Image Analysis, and Statistical Analysis

Oocytes were mounted on a glass slide in SlowFade Gold (Invitrogen) and images were taken with a 63x lens on a Leica SP8 or Stellaris microscope. Protein intensities were quantified using Imaris image analysis software (Bitplane). To measure the intensity of proteins at kinetochores, spots marking the kinetochores and random areas of the background were created. The average intensity of the protein of interest at the background was subtracted from the intensity at each kinetochore to generate the normalized intensity value. GraphPad Prism software was used to graph data and perform statistical analysis. All statistical tests and sample sizes are reported in the figure legends.

### Data availability statement

All the data underlying this study are available in the published article and its online supplementary material.

## Acknowledgements

We thank Marina Druzhinina for technical assistance, Christian Lehner, and Rager Karess for providing Drosophila stocks and antibodies. Stocks obtained from the Bloomington Drosophila Stock Center (NIH P40OD018537) were used in this study. The E7 tubulin monoclonal antibody was obtained from the Developmental Studies Hybridoma Bank, created by the NICHD of the NIH and maintained at The University of Iowa, Department of Biology, Iowa City, IA 52242. Image acquisition and analysis was made possible by the Waksman Institute Shared Imaging Core Facility and The Human Genetics Institute Imaging Core Facility at Rutgers. This work was supported by NIH grant GM101955 to K.S.M.

Figure S 1. **Validation of *rod* and *mps1* reagents.**

(A) Wild-type oocyte with ROD^GFP^ in green, ZW10^HA^ in red, DNA in blue, and tubulin in white. Single channel images show ROD^GFP^ (middle) and ZW10^HA^ (right). (B) ROD^GFP^ localization in wild-type and *rod^RNAi^* oocytes with ROD^GFP^ in green, DNA in blue, CENP-C in white, and tubulin in red. Single channel images (right) show ROD^GFP^. (C) Quantification of ROD^GFP^ intensity at kinetochores, normalized to background GFP signal in wild-type and *rod^RNAi^* oocytes (n = 142 and 126 kinetochores). Error bars show mean ± s.d.; ****P<0.0001 (unpaired two-tailed t test). (D) MPS1^GFP^ localization in wild-type and *Mps1^RNAi^* oocytes with MPS1^GFP^ in green, DNA in blue, CENP-C in white, and tubulin in red. Single channel images (right) show MPS1^GFP^. (E) Quantification of MPS1^GFP^ intensity at kinetochores, normalized to background GFP signal in wild-type and *Mps1^RNAi^* oocytes (n = 75 and 71 kinetochores). Error bars show mean ± s.d.; ****P<0.0001 (unpaired two-tailed t test). All images are maximum intensity projections of z stacks. Scale bars represent 5 µm.

Figure S 2. **MAD1 and INCENP localization depends on SPC105R.**

Control (*Spc105R^B^*) or mutant oocytes were incubated for one hour in 250 µM colchicine. (A) MAD1^GFP^ (green) localization in indicated genotypes, with DNA in blue, CENP-C in white, and tubulin in red. Single channel images (bottom) show MAD1^GFP^. (B) Quantification of MAD1^GFP^ intensity at kinetochores, normalized to background GFP signal (from left to right, n = 91, 256, 224, 174, 212, 208, and 236 kinetochores). Error bars show mean ± s.d.; ****P<0.0001, ***P=0.0006 (unpaired two-tailed t test). (C) INCENP localization (green) in indicated genotypes with DNA in blue, CENP-C in white, and tubulin in red. Single channel images (bottom) show INCENP. (D) Quantification of INCENP presence at kinetochores (from left to right, n = 16, 29, 11, 31, 20, 18, 21, and 11 oocytes). Error bars show mean ± s.d.; ****P<0.0001, *P=0.02 (Fisher’s exact test). All images are maximum intensity projections of z stacks. Scale bars represent 5 µm. All *Spc105R* mutants are in an *Spc105R^RNAi^* background targeting the endogenous *Spc105R*.

Figure S 3. **Structure of SPC105R and mutant variants.**

A schematic of the *Spc105R* mutants used in this study. The coordinates on the schematic represent the first amino acid of each domain. The N-terminal includes SLRK and RISF. Following this is a domain with three MELT-like motifs, a region that contains two KI-like repeats, a central domain containing repeats with the consensus ExxEED, and the C-terminal region containing coiled-coil motifs.

Figure S 4. **MPS1 localization depends on SPC105R and NDC80.**

(A) MPS1^GFP^ localization in wild-type, *Spc105R^RNAi^*, and *Ndc80^RNAi^* oocytes, with MPS1^GFP^ in green, DNA in blue, CENP-C in white, and tubulin in red. Single channel images (bottom) show MPS1^GFP^. (B) Quantification of MPS1^GFP^ intensity at kinetochores, normalized to background GFP signal in indicated oocytes (from left to right, n = 389, 195, 139, 160, 165, 198, 128, and 143 kinetochores). Error bars show mean ± s.d.; ****P<0.0001 (unpaired two-tailed t test). (C) Quantification of MPS1^GFP^ intensity at kinetochores, normalized to background GFP signal in wild-type and *Ndc80^RNAi^* oocytes (n=114 and 148 kinetochores). Error bars show mean ± s.d.; ****P<0.0001 (unpaired two-tailed t test). (D) MPS1^GFP^ localization in the indicated *Spc105R* mutants with MPS1^GFP^ in green, DNA in blue, CENP-C in white, and tubulin in red. All mutants are in an *Spc105R^RNAi^* background targeting the endogenous *Spc105R*. Single channel images (bottom) show MPS1^GFP^. All images are maximum intensity projections of z stacks. Scale bars represent 5 µm.

Figure S 5. **ROD and SPC105R localization in Z*w10-Spc105R^C^* oocytes.**

(A) SPC105R localization in Z*w10-Spc105R^C^*, either in the presence or absence of *Spc105R^RNAi^*. SPC105R (green) was detected using an antibody which recognizes the N-terminal regions of SPC105R and does not detect SPC105R^C^. DNA is in blue, CENP-C in white, and tubulin in red. All images are maximum intensity projections of z stacks. (B) ROD^GFP^ localization in oocytes expressing Z*w10-Spc105R^C^,* either in the presence or absence of *Spc105R^RNAi^*. ROD^GFP^ is in green, DNA in blue, CENP-C in white, and tubulin in red. Scale bars represent 5 µm.

Figure S 6. **Localization of Spindly variants and dependence of Spindly on ROD.**

(A) Examples of normal (left) and abnormal (right) ROD^GFP^ streaming in *Spindly^RNAi^* oocytes, with ROD^GFP^ in green, DNA in blue, CENP-C in white, and tubulin in red. Single channel images (bottom) show ROD^GFP^. Arrow points to abnormal streaming, which is defined as having ROD^GFP^ in the central region between the centromeres. (B) Spindly^GFP^ localization in wild-type or mutants of *Spindly*. Spindly^GFP^ is in green, DNA in blue, CENP-C in white, and tubulin in red. (C) Localization of Spindly^ΔSB.GFP^ in *rod^RNAi^* oocytes, with Spindly^GFP^ in green, DNA in blue, CENP-C in white, and tubulin in red. Spindly^GFP^ and CENP-C are shown below the merged images. All images are maximum intensity projections of z stacks. Scale bars represent 5 µm. (D) Quantification of Spindly^GFP^ intensity at kinetochores in the indicated oocytes (from left to right, n = 123, 164, 83, and 75 kinetochores). Error bars show mean ± s.d.; ****P<0.0001 (unpaired two-tailed t test).

## References

1. McAinsh AD, Kops G. Principles and dynamics of spindle assembly checkpoint signalling. Nat Rev Mol Cell Biol. 2023;24(8):543–59. Epub 20230324. doi: 10.1038/s41580-023-00593-z. PubMed PMID: 36964313.

2. Lara-Gonzalez P, Pines J, Desai A. Spindle assembly checkpoint activation and silencing at kinetochores. Seminars in cell & developmental biology. 2021;117:86–98. Epub 20210629. doi: 10.1016/j.semcdb.2021.06.009. PubMed PMID: 34210579; PubMed Central PMCID: PMCPMC8406419.

3. Krenn V, Musacchio A. The Aurora B Kinase in Chromosome Bi-Orientation and Spindle Checkpoint Signaling. Front Oncol. 2015;5:225. Epub 2015/11/04. doi: 10.3389/fonc.2015.00225. PubMed PMID: 26528436; PubMed Central PMCID: PMCPMC4607871.

4. Barbosa J, Sunkel CE, Conde C. The Role of Mitotic Kinases and the RZZ Complex in Kinetochore-Microtubule Attachments: Doing the Right Link. Front Cell Dev Biol. 2022;10:787294. Epub 20220128. doi: 10.3389/fcell.2022.787294. PubMed PMID: 35155423; PubMed Central PMCID: PMCPMC8832123.

5. Lampson MA, Grishchuk EL. Mechanisms to Avoid and Correct Erroneous Kinetochore-Microtubule Attachments. Biology (Basel). 2017;6(1). Epub 2017/01/10. doi: 10.3390/biology6010001. PubMed PMID: 28067761; PubMed Central PMCID: PMCPMC5371994.

6. Kitajima TS. Mechanisms of kinetochore-microtubule attachment errors in mammalian oocytes. Dev Growth Differ. 2018;60(1):33–43. Epub 2018/01/11. doi: 10.1111/dgd.12410. PubMed PMID: 29318599.

7. Mihajlovic AI, FitzHarris G. Segregating Chromosomes in the Mammalian Oocyte. Curr Biol. 2018;28(16):R895–R907. Epub 2018/08/22. doi: 10.1016/j.cub.2018.06.057. PubMed PMID: 30130515.

8. Webster A, Schuh M. Mechanisms of Aneuploidy in Human Eggs. Trends Cell Biol. 2017;27(1):55–68. Epub 2016/10/25. doi: 10.1016/j.tcb.2016.09.002. PubMed PMID: 27773484.

9. Charalambous C, Webster A, Schuh M. Aneuploidy in mammalian oocytes and the impact of maternal ageing. Nat Rev Mol Cell Biol. 2023;24(1):27–44. Epub 20220906. doi: 10.1038/s41580-022-00517-3. PubMed PMID: 36068367.

10. Touati SA, Wassmann K. How oocytes try to get it right: spindle checkpoint control in meiosis. Chromosoma. 2016;125(2):321–35. Epub 20150811. doi: 10.1007/s00412-015-0536-7. PubMed PMID: 26255654.

11. Wu T, Gu H, Luo Y, Wang L, Sang Q. Meiotic defects in human oocytes: Potential causes and clinical implications. Bioessays. 2022;44(12):e2200135. Epub 20221007. doi: 10.1002/bies.202200135. PubMed PMID: 36207289.

12. Liu D, Shao H, Wang H, Liu XJ. Meiosis I in Xenopus oocytes is not error-prone despite lacking spindle assembly checkpoint. Cell Cycle. 2014;13(10):1602–6. Epub 20140319. doi: 10.4161/cc.28562. PubMed PMID: 24646611; PubMed Central PMCID: PMCPMC4050165.

13. Shao H, Li R, Ma C, Chen E, Liu XJ. Xenopus oocyte meiosis lacks spindle assembly checkpoint control. J Cell Biol. 2013;201(2):191–200. Epub 20130408. doi: 10.1083/jcb.201211041. PubMed PMID: 23569212; PubMed Central PMCID: PMCPMC3628510.

14. Gui L, Homer H. Spindle assembly checkpoint signalling is uncoupled from chromosomal position in mouse oocytes. Development. 2012;139(11):1941–6. PubMed PMID: 22513372.

15. Lane SI, Yun Y, Jones KT. Timing of anaphase-promoting complex activation in mouse oocytes is predicted by microtubule-kinetochore attachment but not by bivalent alignment or tension. Development. 2012;139(11):1947–55. PubMed PMID: 22513370.

16. Kyogoku H, Kitajima TS. The large cytoplasmic volume of oocyte. J Reprod Dev. 2023;69(1):1–9. Epub 20221126. doi: 10.1262/jrd.2022-101. PubMed PMID: 36436912; PubMed Central PMCID: PMCPMC9939283.

17. Radford SJ, Nguyen AL, Schindler K, McKim KS. The chromosomal basis of meiotic acentrosomal spindle assembly and function in oocytes. Chromosoma. 2017;126(3):351–64. Epub 2016/11/12. doi: 10.1007/s00412-016-0618-1. PubMed PMID: 27837282; PubMed Central PMCID: PMCPMC5426991.

18. Blengini CS, Schindler K. Acentriolar spindle assembly in mammalian female meiosis and the consequences of its perturbations on human reproduction†. Biol Reprod. 2022;106(2):253–63. doi: 10.1093/biolre/ioab210. PubMed PMID: 34791041; PubMed Central PMCID: PMCPMC8862719.

19. Wartosch L, Schindler K, Schuh M, Gruhn JR, Hoffmann ER, McCoy RC, et al. Origins and mechanisms leading to aneuploidy in human eggs. Prenat Diagn. 2021;41(5):620–30. Epub 20210322. doi: 10.1002/pd.5927. PubMed PMID: 33860956; PubMed Central PMCID: PMCPMC8237340.

20. Singleton MR. Getting to the heart of an unusual kinetochore. Open Biol. 2016;6(4):160040. doi: 10.1098/rsob.160040. PubMed PMID: 27249344; PubMed Central PMCID: PMCPMC4852463.

21. Radford SJ, Hoang TL, Głuszek AA, Ohkura H, McKim KS. Lateral and End-On Kinetochore Attachments Are Coordinated to Achieve Bi-orientation in Drosophila Oocytes. PLoS Genet. 2015;11(10):e1005605. doi: 10.1371/journal.pgen.1005605. PubMed PMID: 26473960.

22. Joshi JN, Changela N, Mahal L, Jang J, Defosse T, Wang LI, et al. Meiosis-specific functions of kinetochore protein SPC105R required for chromosome segregation in Drosophila oocytes. Mol Biol Cell. 2024:mbcE24020067. Epub 20240612. doi: 10.1091/mbc.E24-02-0067. PubMed PMID: 38865189.

23. Cheerambathur DK, Gassmann R, Cook B, Oegema K, Desai A. Crosstalk between microtubule attachment complexes ensures accurate chromosome segregation. Science. 2013;342(6163):1239-42. Epub 2013/11/16. doi: 10.1126/science.1246232. PubMed PMID: 24231804; PubMed Central PMCID: PMCPMC3885540.

24. Barbosa J, Martins T, Bange T, Tao L, Conde C, Sunkel C. Polo regulates Spindly to prevent premature stabilization of kinetochore-microtubule attachments. EMBO J. 2020;39(2):e100789. Epub 2019/12/19. doi: 10.15252/embj.2018100789. PubMed PMID: 31849090; PubMed Central PMCID: PMCPMC6960449.

25. Basto R, Scaerou F, Mische S, Wojcik E, Lefebvre C, Gomes R, et al. In vivo dynamics of the rough deal checkpoint protein during Drosophila mitosis. Curr Biol. 2004;14(1):56–61. PubMed PMID: 14711415.

26. Williams BC, Gatti M, Goldberg ML. Bipolar spindle attachments affect redistributions of ZW10, a Drosophila centromere/kinetochore component required for accurate chromosome segregation. J Cell Biol. 1996;134(5):1127–40. doi: 10.1083/jcb.134.5.1127. PubMed PMID: 8794856; PubMed Central PMCID: PMCPMC2120981.

27. Scaërou F, Starr DA, Piano F, Papoulas O, Karess RE, Goldberg ML. The ZW10 and Rough Deal checkpoint proteins function together in a large, evolutionarily conserved complex targeted to the kinetochore. J Cell Sci. 2001;114(Pt 17):3103–14. doi: 10.1242/jcs.114.17.3103. PubMed PMID: 11590237.

28. Kops G, Gassmann R. Crowning the Kinetochore: The Fibrous Corona in Chromosome Segregation. Trends Cell Biol. 2020;30(8):653–67. Epub 2020/05/11. doi: 10.1016/j.tcb.2020.04.006. PubMed PMID: 32386879.

29. Pereira C, Reis RM, Gama JB, Celestino R, Cheerambathur DK, Carvalho AX, et al. Self-Assembly of the RZZ Complex into Filaments Drives Kinetochore Expansion in the Absence of Microtubule Attachment. Current biology : CB. 2018;28(21):3408–21.e8. Epub 2018/11/13. doi: 10.1016/j.cub.2018.08.056. PubMed PMID: 30415699; PubMed Central PMCID: PMCPMC6224608.

30. Marston AL, Wassmann K. Multiple Duties for Spindle Assembly Checkpoint Kinases in Meiosis. Front Cell Dev Biol. 2017;5:109. Epub 20171213. doi: 10.3389/fcell.2017.00109. PubMed PMID: 29322045; PubMed Central PMCID: PMCPMC5733479.

31. Wang LI, DeFosse T, Jang JK, Battaglia RA, Wagner VF, McKim KS. Borealin directs recruitment of the CPC to oocyte chromosomes and movement to the microtubules. J Cell Biol. 2021;220(6). Epub 2021/04/10. doi: 10.1083/jcb.202006018. PubMed PMID: 33836043.

32. Radford SJ, Jang JK, McKim KS. The Chromosomal Passenger Complex is required for Meiotic Acentrosomal Spindle Assembly and Chromosome Bi-orientation. Genetics. 2012;192:417–29. PubMed PMID: 22865736.

33. McKim KS, Jang JK, Theurkauf WE, Hawley RS. Mechanical basis of meiotic metaphase arrest. Nature. 1993;362(6418):364-6. PubMed PMID: 8455723.

34. Saurin AT. Kinase and Phosphatase Cross-Talk at the Kinetochore. Front Cell Dev Biol. 2018;6:62. Epub 2018/07/05. doi: 10.3389/fcell.2018.00062. PubMed PMID: 29971233; PubMed Central PMCID: PMCPMC6018199.

35. Caldas GV, DeLuca JG. KNL1: bringing order to the kinetochore. Chromosoma. 2014;123(3):169–81. doi: 10.1007/s00412-013-0446-5. PubMed PMID: 24310619; PubMed Central PMCID: PMCPMC4032621.

36. Hayward D, Alfonso-Pérez T, Cundell MJ, Hopkins M, Holder J, Bancroft J, et al. CDK1-CCNB1 creates a spindle checkpoint-permissive state by enabling MPS1 kinetochore localization. J Cell Biol. 2019;218(4):1182–99. Epub 20190123. doi: 10.1083/jcb.201808014. PubMed PMID: 30674582; PubMed Central PMCID: PMCPMC6446832.

37. Gilliland WD, Hughes SE, Cotitta JL, Takeo S, Xiang Y, Hawley RS. The multiple roles of mps1 in Drosophila female meiosis. PLoS Genet. 2007;3(7):e113. doi: 10.1371/journal.pgen.0030113. PubMed PMID: 17630834; PubMed Central PMCID: PMCPMC1914070.

38. Gilliland WD, Wayson SM, Hawley RS. The meiotic defects of mutants in the Drosophila mps1 gene reveal a critical role of Mps1 in the segregation of achiasmate homologs. Curr Biol. 2005;15(7):672–7. doi: 10.1016/j.cub.2005.02.062. PubMed PMID: 15823541.

39. Bourouh M, Dhaliwal R, Rana K, Sinha S, Guo Z, Swan A. Distinct and Overlapping Requirements for Cyclins A, B and B3 in Drosophila Female Meiosis. G3 (Bethesda, Md. 2016. doi: 10.1534/g3.116.033050. PubMed PMID: 27652889; PubMed Central PMCID: PMCPMC5100870.

40. Malmanche N, Owen S, Gegick S, Steffensen S, Tomkiel JE, Sunkel CE. Drosophila BubR1 is essential for meiotic sister-chromatid cohesion and maintenance of synaptonemal complex. Curr Biol. 2007;17(17):1489–97. PubMed PMID: 17702574.

41. Buffin E, Emre D, Karess RE. Flies without a spindle checkpoint. Nat Cell Biol. 2007;9(5):565–72. doi: 10.1038/ncb1570. PubMed PMID: 17417628.

42. Emre D, Terracol R, Poncet A, Rahmani Z, Karess RE. A mitotic role for Mad1 beyond the spindle checkpoint. J Cell Sci. 2011;124(Pt 10):1664–71. Epub 20110421. doi: 10.1242/jcs.081216. PubMed PMID: 21511728.

43. Hughes SE, Miller DE, Miller AL, Hawley RS. Female Meiosis: Synapsis, Recombination, and Segregation in Drosophila melanogaster. Genetics. 2018;208(3):875–908. Epub 2018/03/01. doi: 10.1534/genetics.117.300081. PubMed PMID: 29487146; PubMed Central PMCID: PMCPMC5844340.

44. Theurkauf WE, Hawley RS. Meiotic spindle assembly in Drosophila females: behavior of nonexchange chromosomes and the effects of mutations in the nod kinesin-like protein. J Cell Biol. 1992;116(5):1167–80. PubMed PMID: 1740471.

45. Costa MFA, Ohkura H. The molecular architecture of the meiotic spindle is remodeled during metaphase arrest in oocytes. J Cell Biol. 2019;218(9):2854–64. Epub 2019/07/07. doi: 10.1083/jcb.201902110. PubMed PMID: 31278080; PubMed Central PMCID: PMCPMC6719438.

46. Carpenter AT, Sandler L. On recombination-defective meiotic mutants in Drosophila melanogaster. Genetics. 1974;76(3):453–75. PubMed PMID: 4208856.

47. Jang JK, Gladstein AC, Das A, Shapiro JG, Sisco ZL, McKim KS. Multiple pools of PP2A regulate spindle assembly, kinetochore attachments, and cohesion in Drosophila oocytes. J Cell Sci. 2021;134(14):jcs254037. Epub 2021/06/24. doi: 10.1242/jcs.254037. PubMed PMID: 34160620.

48. Wu J, Larreategui-Aparicio A, Lambers MLA, Bodor DL, Klaasen SJ, Tollenaar E, et al. Microtubule nucleation from the fibrous corona by LIC1-pericentrin promotes chromosome congression. Curr Biol. 2023;33(5):912–25 e6. Epub 20230130. doi: 10.1016/j.cub.2023.01.010. PubMed PMID: 36720222; PubMed Central PMCID: PMCPMC10017265.

49. Menant A, Karess RE. Mutations in the Drosophila rough deal gene affecting RZZ kinetochore function. Biol Cell. 2020;112(10):300–15. Epub 20200720. doi: 10.1111/boc.201900105. PubMed PMID: 32602944.

50. Wojcik E, Basto R, Serr M, Scaërou F, Karess R, Hays T. Kinetochore dynein: its dynamics and role in the transport of the Rough deal checkpoint protein. Nature cell biology. 2001;3(11):1001–7. Epub 2001/11/21. doi: 10.1038/ncb1101-1001. PubMed PMID: 11715021.

51. Griffis ER, Stuurman N, Vale RD. Spindly, a novel protein essential for silencing the spindle assembly checkpoint, recruits dynein to the kinetochore. J Cell Biol. 2007;177(6):1005-15. doi: 10.1083/jcb.200702062. PubMed PMID: 17576797; PubMed Central PMCID: PMCPMC2064361.

52. Wainman A, Creque J, Williams B, Williams EV, Bonaccorsi S, Gatti M, et al. Roles of the Drosophila NudE protein in kinetochore function and centrosome migration. J Cell Sci. 2009;122(Pt 11):1747–58. Epub 2009/05/07. doi: 10.1242/jcs.041798. PubMed PMID: 19417004; PubMed Central PMCID: PMCPMC2684831.

53. Gassmann R, Essex A, Hu JS, Maddox PS, Motegi F, Sugimoto A, et al. A new mechanism controlling kinetochore-microtubule interactions revealed by comparison of two dynein-targeting components: SPDL-1 and the Rod/Zwilch/Zw10 complex. Genes Dev. 2008;22(17):2385–99. doi: 10.1101/gad.1687508. PubMed PMID: 18765790; PubMed Central PMCID: PMCPMC2532926.

54. Clemente GD, Hannaford MR, Beati H, Kapp K, Januschke J, Griffis ER, et al. Requirement of the Dynein-Adaptor Spindly for Mitotic and Post-Mitotic Functions in Drosophila. J Dev Biol. 2018;6(2). Epub 2018/04/05. doi: 10.3390/jdb6020009. PubMed PMID: 29615558; PubMed Central PMCID: PMCPMC6027351.

55. Davis-Roca AC, Muscat CC, Wignall SM. Caenorhabditis elegans oocytes detect meiotic errors in the absence of canonical end-on kinetochore attachments. J Cell Biol. 2017;216(5):1243–53. Epub 20170329. doi: 10.1083/jcb.201608042. PubMed PMID: 28356326; PubMed Central PMCID: PMCPMC5412562.

56. Jones KT, Lane SI. Molecular causes of aneuploidy in mammalian eggs. Development. 2013;140(18):3719–30. doi: 10.1242/dev.090589. PubMed PMID: 23981655.

57. Touati SA, Buffin E, Cladiere D, Hached K, Rachez C, van Deursen JM, et al. Mouse oocytes depend on BubR1 for proper chromosome segregation but not for prophase I arrest. Nat Commun. 2015;6:6946. Epub 2015/04/22. doi: 10.1038/ncomms7946. PubMed PMID: 25897860; PubMed Central PMCID: PMCPMC4439927.

58. Qiao JY, Zhou Q, Xu K, Yue W, Lei WL, Li YY, et al. Mad2 is dispensable for accurate chromosome segregation but becomes essential when oocytes are subjected to environmental stress. Development. 2023;150(14). Epub 20230724. doi: 10.1242/dev.201398. PubMed PMID: 37485540.

59. Batiha O, Swan A. Evidence that the spindle assembly checkpoint does not regulate APC(Fzy) activity in Drosophila female meiosis. Genome. 2012;55(1):63–7. Epub 20111223. doi: 10.1139/g11-079. PubMed PMID: 22196012.

60. Jang JK, Messina L, Erdman MB, Arbel T, Hawley RS. Induction of metaphase arrest in Drosophila oocytes by chiasma-based kinetochore tension. Science. 1995;268(5219):1917-9.

61. !!! INVALID CITATION !!! [22, 46].

62. !!! INVALID CITATION !!! [31, 58].

63. !!! INVALID CITATION !!! [60].

64. Silió V, McAinsh AD, Millar JB. KNL1-Bubs and RZZ Provide Two Separable Pathways for Checkpoint Activation at Human Kinetochores. Dev Cell. 2015;35(5):600–13. Epub 2015/12/15. doi: 10.1016/j.devcel.2015.11.012. PubMed PMID: 26651294.

65. Pauleau AL, Bergner A, Kajtez J, Erhardt S. The checkpoint protein Zw10 connects CAL1-dependent CENP-A centromeric loading and mitosis duration in Drosophila cells. PLoS Genet. 2019;15(9):e1008380. Epub 20190925. doi: 10.1371/journal.pgen.1008380. PubMed PMID: 31553715; PubMed Central PMCID: PMCPMC6779278.

66. O’Tousa J. Meiotic chromosome behavior influenced by mutation-altered disjunction in Drosophila melanogaster females. Genetics. 1982;102:503–24.

67. Hached K, Xie SZ, Buffin E, Cladiere D, Rachez C, Sacras M, et al. Mps1 at kinetochores is essential for female mouse meiosis I. Development. 2011;138(11):2261–71. PubMed PMID: 21558374.

68. Sarangapani KK, Koch LB, Nelson CR, Asbury CL, Biggins S. Kinetochore-bound Mps1 regulates kinetochore-microtubule attachments via Ndc80 phosphorylation. J Cell Biol. 2021;220(12). Epub 20211014. doi: 10.1083/jcb.202106130. PubMed PMID: 34647959; PubMed Central PMCID: PMCPMC8641409.

69. Hiruma Y, Sacristan C, Pachis ST, Adamopoulos A, Kuijt T, Ubbink M, et al. CELL DIVISION CYCLE. Competition between MPS1 and microtubules at kinetochores regulates spindle checkpoint signaling. Science. 2015;348(6240):1264-7. Epub 20150611. doi: 10.1126/science.aaa4055. PubMed PMID: 26068855.

70. Ji Z, Gao H, Yu H. CELL DIVISION CYCLE. Kinetochore attachment sensed by competitive Mps1 and microtubule binding to Ndc80C. Science. 2015;348(6240):1260-4. doi: 10.1126/science.aaa4029. PubMed PMID: 26068854.

71. Kemmler S, Stach M, Knapp M, Ortiz J, Pfannstiel J, Ruppert T, et al. Mimicking Ndc80 phosphorylation triggers spindle assembly checkpoint signalling. EMBO J. 2009;28(8):1099–110. Epub 20090319. doi: 10.1038/emboj.2009.62. PubMed PMID: 19300438; PubMed Central PMCID: PMCPMC2683709.

72. Hayward D, Roberts E, Gruneberg U. MPS1 localizes to end-on microtubule-attached kinetochores to promote microtubule release. Curr Biol. 2022;32(23):5200–8 e8. Epub 20221116. doi: 10.1016/j.cub.2022.10.047. PubMed PMID: 36395767.

73. Buffin E, Lefebvre C, Huang J, Gagou ME, Karess RE. Recruitment of Mad2 to the kinetochore requires the Rod/Zw10 complex. Curr Biol. 2005;15(9):856–61. PubMed PMID: 15886105.

74. Kops GJ, Kim Y, Weaver BA, Mao Y, McLeod I, Yates JR, 3rd, et al. ZW10 links mitotic checkpoint signaling to the structural kinetochore. J Cell Biol. 2005;169(1):49–60. doi: 10.1083/jcb.200411118. PubMed PMID: 15824131; PubMed Central PMCID: PMCPMC1351127.

75. Tromer E, Snel B, Kops GJ. Widespread Recurrent Patterns of Rapid Repeat Evolution in the Kinetochore Scaffold KNL1. Genome Biol Evol. 2015;7(8):2383–93. Epub 2015/08/09. doi: 10.1093/gbe/evv140. PubMed PMID: 26254484; PubMed Central PMCID: PMCPMC4558858.

76. Caldas GV, Lynch TR, Anderson R, Afreen S, Varma D, DeLuca JG. The RZZ complex requires the N-terminus of KNL1 to mediate optimal Mad1 kinetochore localization in human cells. Open Biol. 2015;5(11). doi: 10.1098/rsob.150160. PubMed PMID: 26581576; PubMed Central PMCID: PMCPMC4680571.

77. Zhang G, Lischetti T, Hayward DG, Nilsson J. Distinct domains in Bub1 localize RZZ and BubR1 to kinetochores to regulate the checkpoint. Nat Commun. 2015;6:7162. Epub 20150602. doi: 10.1038/ncomms8162. PubMed PMID: 26031201; PubMed Central PMCID: PMCPMC4458899.

78. McGory JM, Verma V, Barcelos DM, Maresca TJ. Multimerization of a disordered kinetochore protein promotes accurate chromosome segregation by localizing a core dynein module. J Cell Biol. 2024;223(3). Epub 20240105. doi: 10.1083/jcb.202211122. PubMed PMID: 38180477; PubMed Central PMCID: PMCPMC10770731.

79. Davydenko O, Schultz RM, Lampson MA. Increased CDK1 activity determines the timing of kinetochore-microtubule attachments in meiosis I. J Cell Biol. 2013;202(2):221–9. doi: 10.1083/jcb.201303019. PubMed PMID: 23857768; PubMed Central PMCID: PMCPMC3718970.

80. Kabeche L, Compton DA. Cyclin A regulates kinetochore microtubules to promote faithful chromosome segregation. Nature. 2013;502(7469):110-3. doi: 10.1038/nature12507. PubMed PMID: 24013174; PubMed Central PMCID: PMCPMC3791168.

81. Amin MA, McKenney RJ, Varma D. Antagonism between the dynein and Ndc80 complexes at kinetochores controls the stability of kinetochore-microtubule attachments during mitosis. J Biol Chem. 2018;293(16):5755–65. Epub 2018/02/25. doi: 10.1074/jbc.RA117.001699. PubMed PMID: 29475948; PubMed Central PMCID: PMCPMC5912454.

82. Sacristan C, Ahmad MUD, Keller J, Fermie J, Groenewold V, Tromer E, et al. Dynamic kinetochore size regulation promotes microtubule capture and chromosome biorientation in mitosis. Nat Cell Biol. 2018;20(7):800–10. Epub 2018/06/20. doi: 10.1038/s41556-018-0130-3. PubMed PMID: 29915359; PubMed Central PMCID: PMCPMC6485389.

83. Wang LI, Das A, McKim KS. Sister centromere fusion during meiosis I depends on maintaining cohesins and destabilizing microtubule attachments. PLoS Genet. 2019;15(5):e1008072. Epub 2019/06/01. doi: 10.1371/journal.pgen.1008072. PubMed PMID: 31150390; PubMed Central PMCID: PMCPMC6581285.

84. Barbosa J, Conde C, Sunkel C. RZZ-SPINDLY-DYNEIN: you got to keep ’em separated. Cell Cycle. 2020;19(14):1716–26. Epub 20200616. doi: 10.1080/15384101.2020.1780382. PubMed PMID: 32544383; PubMed Central PMCID: PMCPMC7469663.

85. Barisic M, Sohm B, Mikolcevic P, Wandke C, Rauch V, Ringer T, et al. Spindly/CCDC99 is required for efficient chromosome congression and mitotic checkpoint regulation. Mol Biol Cell. 2010;21(12):1968–81. Epub 2010/04/30. doi: 10.1091/mbc.E09-04-0356. PubMed PMID: 20427577; PubMed Central PMCID: PMCPMC2883941.

86. Gassmann R, Holland AJ, Varma D, Wan X, Civril F, Cleveland DW, et al. Removal of Spindly from microtubule-attached kinetochores controls spindle checkpoint silencing in human cells. Genes & development. 2010;24(9):957–71. Epub 2010/05/05. doi: 10.1101/gad.1886810. PubMed PMID: 20439434; PubMed Central PMCID: PMCPMC2861194.

87. Wu J, Raas MWD, Alcaraz PS, Vos HR, Tromer EC, Snel B, et al. A farnesyl-dependent structural role for CENP-E in expansion of the fibrous corona. J Cell Biol. 2024;223(1). Epub 20231107. doi: 10.1083/jcb.202303007. PubMed PMID: 37934467; PubMed Central PMCID: PMCPMC10630089.

88. McGory JM, Verma V, Barcelos DM, Maresca TJ. Multimerization of a disordered kinetochore protein promotes accurate chromosome segregation by localizing a core dynein module. J Cell Biol. 2024;223(3):e202211122. Epub 20240105. doi: 10.1083/jcb.202211122. PubMed PMID: 38180477; PubMed Central PMCID: PMCPMC10770731.

89. Cmentowski V, Ciossani G, d’Amico E, Wohlgemuth S, Owa M, Dynlacht B, et al. RZZ-Spindly and CENP-E form an integrated platform to recruit dynein to the kinetochore corona. Embo j. 2023;42(24):e114838. Epub 20231120. doi: 10.15252/embj.2023114838. PubMed PMID: 37984321; PubMed Central PMCID: PMCPMC10711656.

90. Althoff F, Karess RE, Lehner CF. Spindle checkpoint-independent inhibition of mitotic chromosome segregation by Drosophila Mps1. Mol Biol Cell. 2012;23(12):2275–91. Epub 20120502. doi: 10.1091/mbc.E12-02-0117. PubMed PMID: 22553353; PubMed Central PMCID: PMCPMC3374747.

91. Fischer MG, Heeger S, Hacker U, Lehner CF. The mitotic arrest in response to hypoxia and of polar bodies during early embryogenesis requires Drosophila Mps1. Curr Biol. 2004;14(22):2019–24. doi: 10.1016/j.cub.2004.11.008. PubMed PMID: 15556864.

92. Zeng Y, Li H, Schweppe NM, Hawley RS, Gilliland WD. Statistical analysis of nondisjunction assays in Drosophila. Genetics. 2010;186(2):505–13. doi: 10.1534/genetics.110.118778. PubMed PMID: 20660647; PubMed Central PMCID: PMCPMC2954469.

93. Défachelles L, Hainline SG, Menant A, Lee LA, Karess RE. A maternal effect rough deal mutation suggests that multiple pathways regulate Drosophila RZZ kinetochore recruitment. J Cell Sci. 2015;128(6):1204–16. Epub 20150122. doi: 10.1242/jcs.165712. PubMed PMID: 25616898; PubMed Central PMCID: PMCPMC4359924.

94. Radford SJ, McKim KS. Techniques for Imaging Prometaphase and Metaphase of Meiosis I in Fixed Drosophila Oocytes. Journal of visualized experiments : JoVE. 2016;116(116):e54666. doi: 10.3791/54666. PubMed PMID: 27842371.

95. Chu DT, Klymkowsky MW. The appearance of acetylated alpha-tubulin during early development and cellular differentiation in Xenopus. Dev Biol. 1989;136(1):104–17. doi: 10.1016/0012-1606(89)90134-6. PubMed PMID: 2680681.

96. Wu C, Singaram V, McKim KS. mei-38 is required for chromosome segregation during meiosis in Drosophila females. Genetics. 2008;180(1):61–72. PubMed PMID: 18757915.

97. Fellmeth JE, Jang JK, Persaud M, Sturm H, Changela N, Parikh A, et al. A dynamic population of prophase CENP-C is required for meiotic chromosome segregation. PLoS Genet. 2023;19(11):e1011066. Epub 20231129. doi: 10.1371/journal.pgen.1011066. PubMed PMID: 38019881.

98. Schittenhelm RB, Chaleckis R, Lehner CF. Intrakinetochore localization and essential functional domains of Drosophila Spc105. EMBO J. 2009;28(16):2374–86. doi: 10.1038/emboj.2009.188. PubMed PMID: 19590494; PubMed Central PMCID: PMCPMC2735171.

99. Sugimura I, Lilly MA. Bruno inhibits the expression of mitotic cyclins during the prophase I meiotic arrest of Drosophila oocytes. Dev Cell. 2006;10(1):127–35. PubMed PMID: 16399084.

100. Schittenhelm RB, Heeger S, Althoff F, Walter A, Heidmann S, Mechtler K, et al. Spatial organization of a ubiquitous eukaryotic kinetochore protein network in Drosophila chromosomes. Chromosoma. 2007;116(4):385–402. PubMed PMID: 17333235.

101. Ni JQ, Zhou R, Czech B, Liu LP, Holderbaum L, Yang-Zhou D, et al. A genome-scale shRNA resource for transgenic RNAi in Drosophila. Nature methods. 2011;8(5):405–7. PubMed PMID: 21460824.

102. Liu H, Jang JK, Graham J, Nycz K, McKim KS. Two genes required for meiotic recombination in Drosophila are expressed from a dicistronic message. Genetics. 2000;154(4):1735–46.

