## Supplemental Figures and Tables for "Distinct checkpoint and homolog biorientation pathways regulate meiosis I in *Drosophila* oocytes"

Table S1

| Genotype | XX | XY | XXY | XO | % NDJ |
|---|---|---|---|---|---|
| *w*$^{1118}$*/matα* | 351 | 317 | 0 | 1 | 0.30 |
| *w*$^{1118}$*/rod*$^{GFP}$*matα* | 363 | 389 | 0 | 1 | 0.27 |
| *w*$^{1118}$*/mps1*$^{GFP}$*matα* | 172 | 149 | 0 | 0 | 0.00 |

Figure S1:

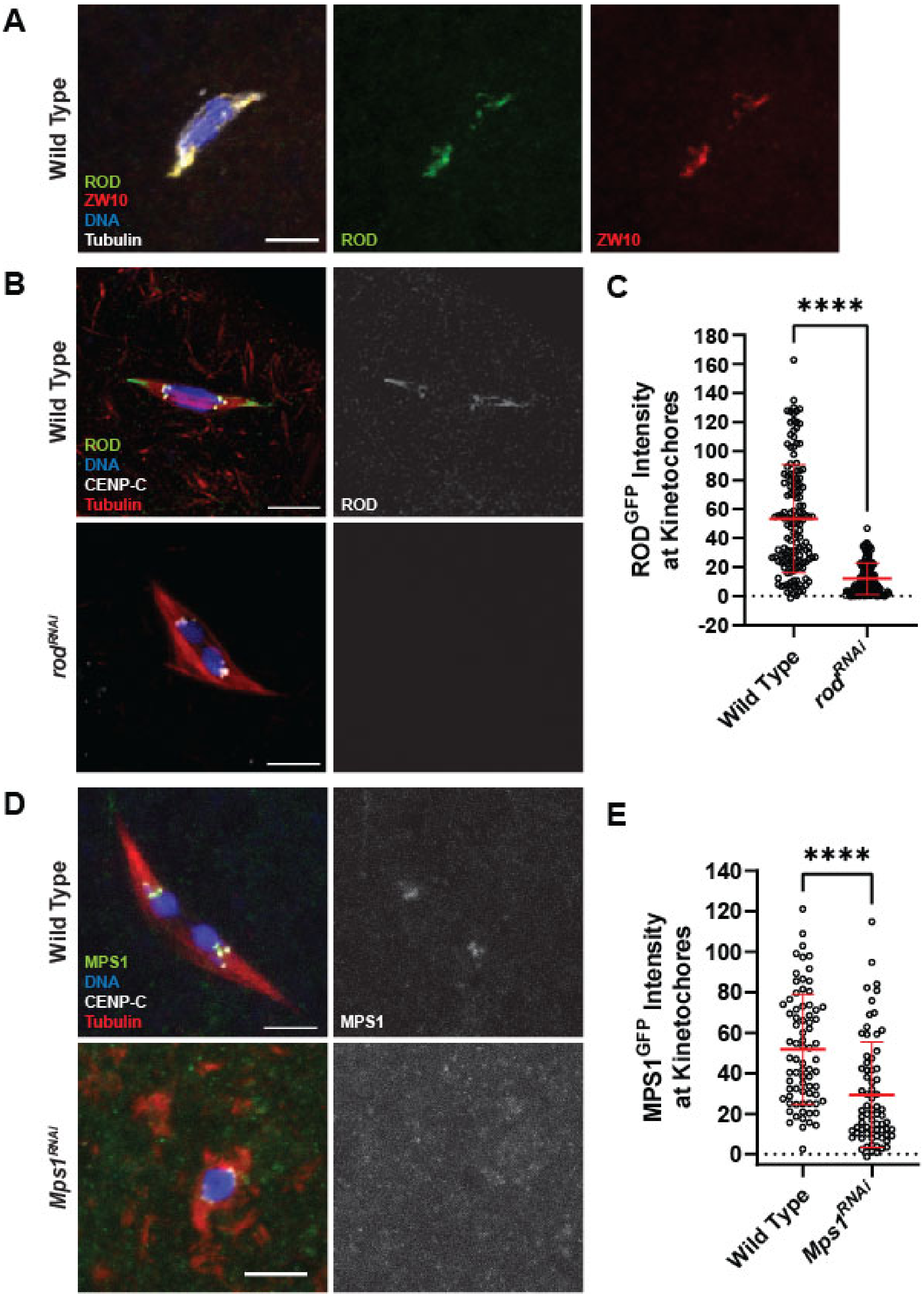

A
Wild Type
ROD
ZW10
DNA
Tubulin
ROD
ZW10
B
Wild Type
ROD
DNA
CENP-C
Tubulin
ROD
rod^RNAi
C
****
ROD^GFP Intensity at Kinetochores
180
160
140
120
100
80
60
40
20
0
-20
Wild Type
rod^RNAi
D
Wild Type
MPS1
DNA
CENP-C
Tubulin
MPS1
Mps1^RNAi
E
****
MPS1^GFP Intensity at Kinetochores
140
120
100
80
60
40
20
0
Wild Type
Mps1^RNAi

Figure S 1. **Validation of *rod* and *mps1* reagents.**

(A) Wild-type oocyte with $ROD^{GFP}$ in green, $ZW10^{HA}$ in red, DNA in blue, and tubulin in white. Single channel images show $ROD^{GFP}$ (middle) and $ZW10^{HA}$ (right). (B) $ROD^{GFP}$ localization in wild-type and $rod^{RNAi}$ oocytes with $ROD^{GFP}$ in green, DNA in blue, CENP-C in white, and tubulin in red. Single channel images (right) show $ROD^{GFP}$. (C) Quantification of $ROD^{GFP}$ intensity at kinetochores, normalized to background GFP signal in wild-type and $rod^{RNAi}$ oocytes (n = 142 and 126 kinetochores). Error bars show mean ± s.d.; ****P<0.0001 (unpaired two-tailed t test). (D) $MPS1^{GFP}$ localization in wild-type and $Mps1^{RNAi}$ oocytes with $MPS1^{GFP}$ in green, DNA in blue, CENP-C in white, and tubulin in red. Single channel images (right) show $MPS1^{GFP}$. (E) Quantification of $MPS1^{GFP}$ intensity at kinetochores, normalized to background GFP signal in wild-type and $Mps1^{RNAi}$ oocytes (n = 75 and 71 kinetochores). Error bars show mean ± s.d.; ****P<0.0001 (unpaired two-tailed t test). All images are maximum intensity projections of z stacks. Scale bars represent 5 μm.

Figure S2

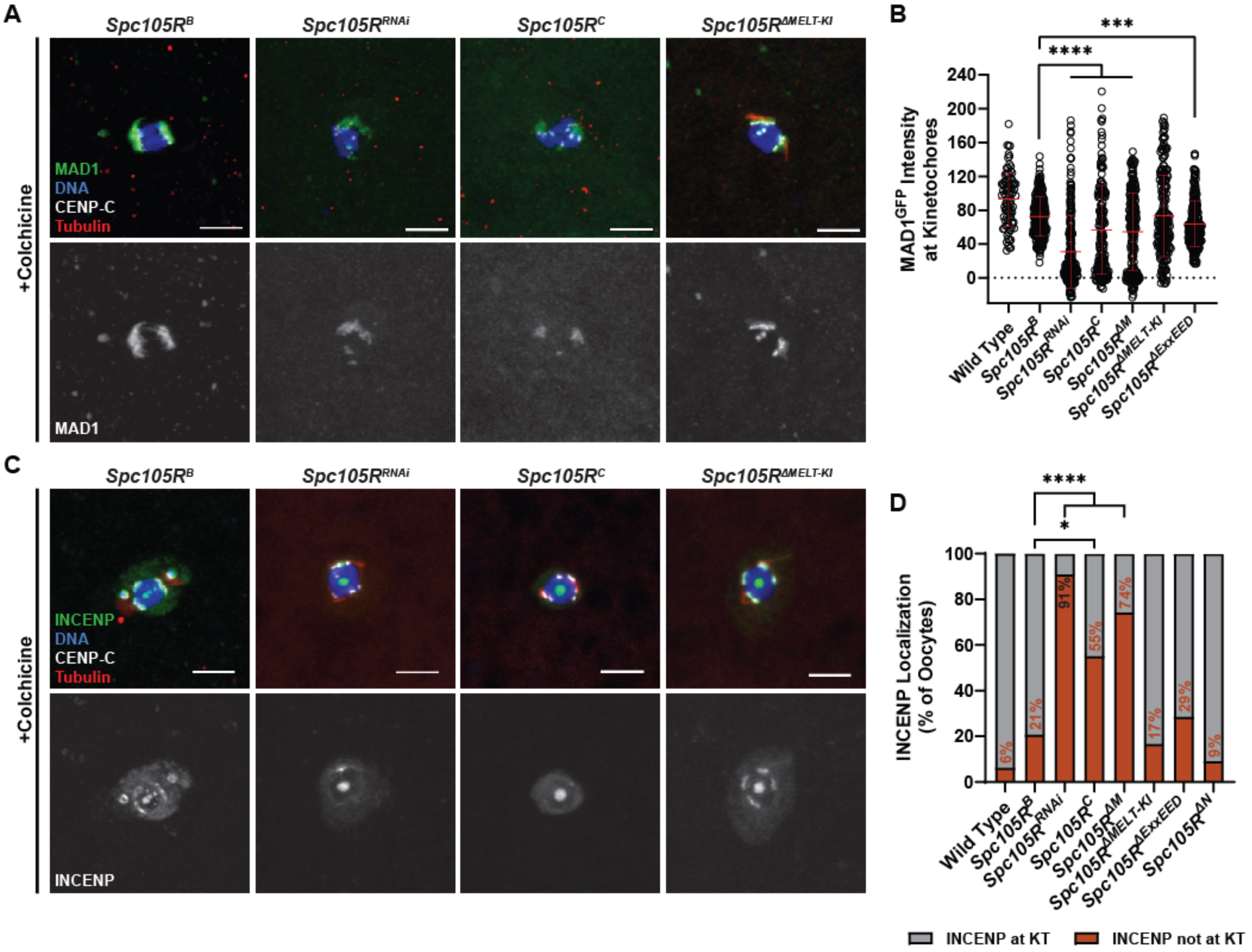


Figure S 2. **MAD1 and INCENP localization depends on SPC105R.**

Control (*Spc105R*$^{B}$) or mutant oocytes were incubated for one hour in 250 µM colchicine. (A) MAD1$^{GFP}$ (green) localization in indicated genotypes, with DNA in blue, CENP-C in white, and tubulin in red. Single channel images (bottom) show MAD1$^{GFP}$. (B) Quantification of MAD1$^{GFP}$ intensity at kinetochores, normalized to background GFP signal (from left to right, n = 91, 256, 224, 174, 212, 208, and 236 kinetochores). Error bars show mean ± s.d.; ****P<0.0001, ***P=0.0006 (unpaired two-tailed t test). (C) INCENP localization (green) in indicated genotypes with DNA in blue, CENP-C in white, and tubulin in red. Single channel images (bottom) show INCENP. (D) Quantification of INCENP presence at kinetochores (from left to right, n = 16, 29, 11, 31, 20, 18, 21, and 11 oocytes). Error bars show mean ± s.d.; ****P<0.0001, *P=0.02 (Fisher's exact test). All images are maximum intensity projections of z stacks. Scale bars represent 5 µm. All *Spc105R* mutants are in an *Spc105R*$^{RNAi}$ background targeting the endogenous *Spc105R*.

Figure S3

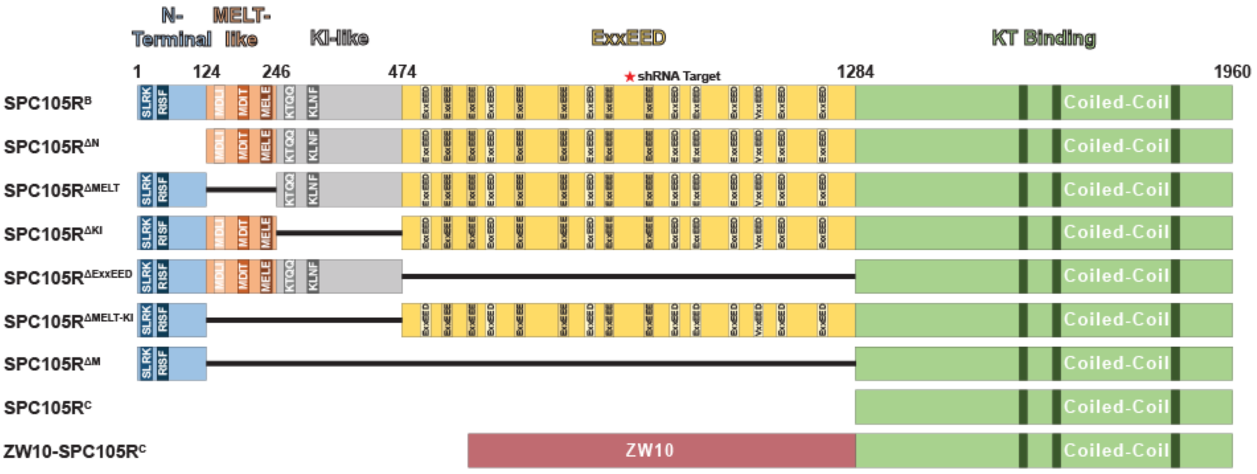


Figure S 3. **Structure of SPC105R and mutant variants.**

A schematic of the *Spc105R* mutants used in this study. The coordinates on the schematic represent the first amino acid of each domain. The N-terminal includes SLRK and RISF. Following this is a domain with three MELT-like motifs, a region that contains two KI-like repeats, a central domain containing repeats with the consensus ExxEED, and the C-terminal region containing coiled-coil motifs.

Figure S4

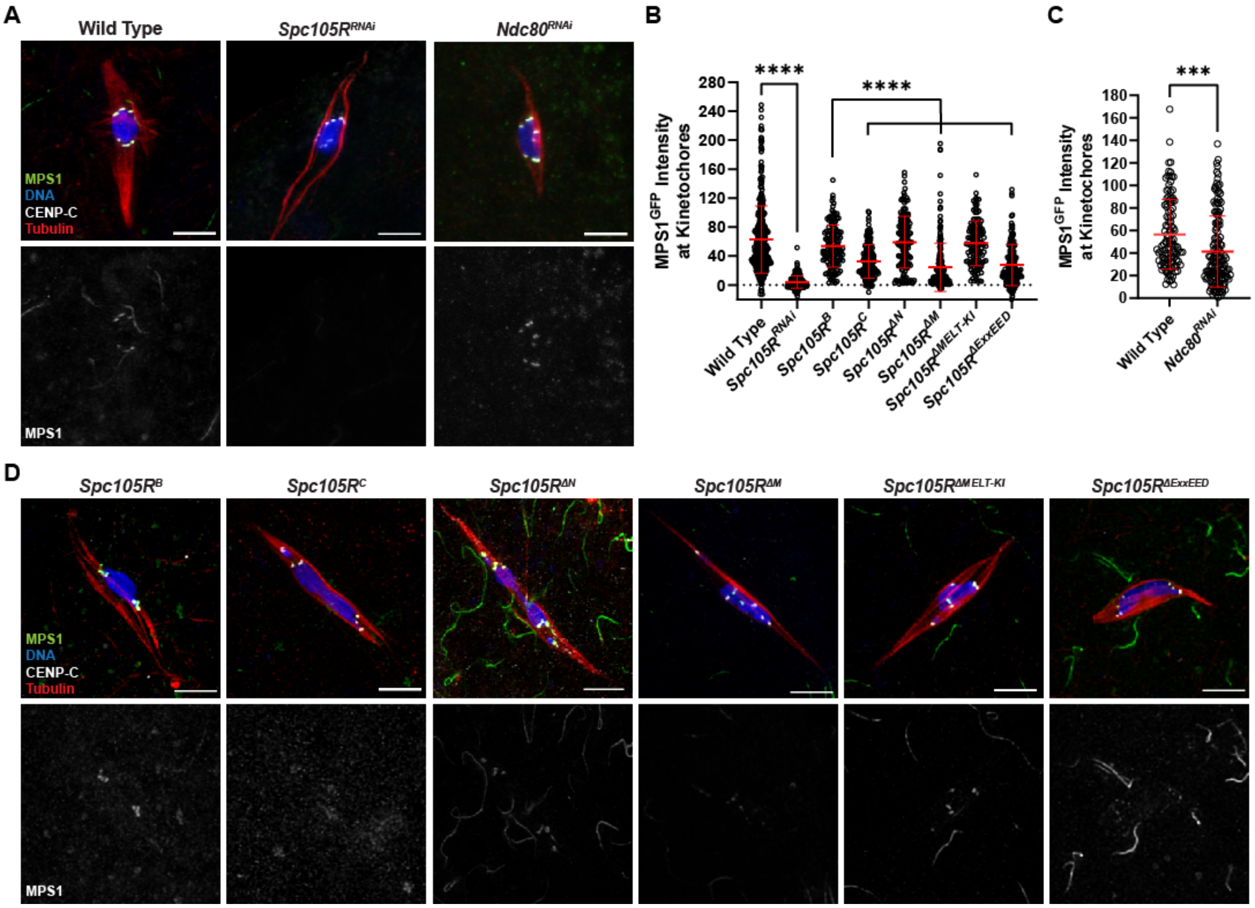


Figure S 4. **MPS1 localization depends on SPC105R and NDC80.**

(A) $MPS1^{GFP}$ localization in wild-type, *Spc105R*$^{RNAi}$, and *Ndc80*$^{RNAi}$ oocytes, with $MPS1^{GFP}$ in green, DNA in blue, CENP-C in white, and tubulin in red. Single channel images (bottom) show $MPS1^{GFP}$. (B) Quantification of $MPS1^{GFP}$ intensity at kinetochores, normalized to background GFP signal in indicated oocytes (from left to right, n = 389, 195, 139, 160, 165, 198, 128, and 143 kinetochores). Error bars show mean ± s.d.; ****P<0.0001 (unpaired two-tailed t test). (C) Quantification of $MPS1^{GFP}$ intensity at kinetochores, normalized to background GFP signal in wild-type and *Ndc80*$^{RNAi}$ oocytes (n=114 and 148 kinetochores). Error bars show mean ± s.d.; ****P<0.0001 (unpaired two-tailed t test). (D) $MPS1^{GFP}$ localization in the indicated *Spc105R* mutants with $MPS1^{GFP}$ in green, DNA in blue, CENP-C in white, and tubulin in red. All mutants are in an *Spc105R*$^{RNAi}$ background targeting the endogenous *Spc105R*. Single channel images (bottom) show $MPS1^{GFP}$. All images are maximum intensity projections of z stacks. Scale bars represent 5 µm.

Figure S5

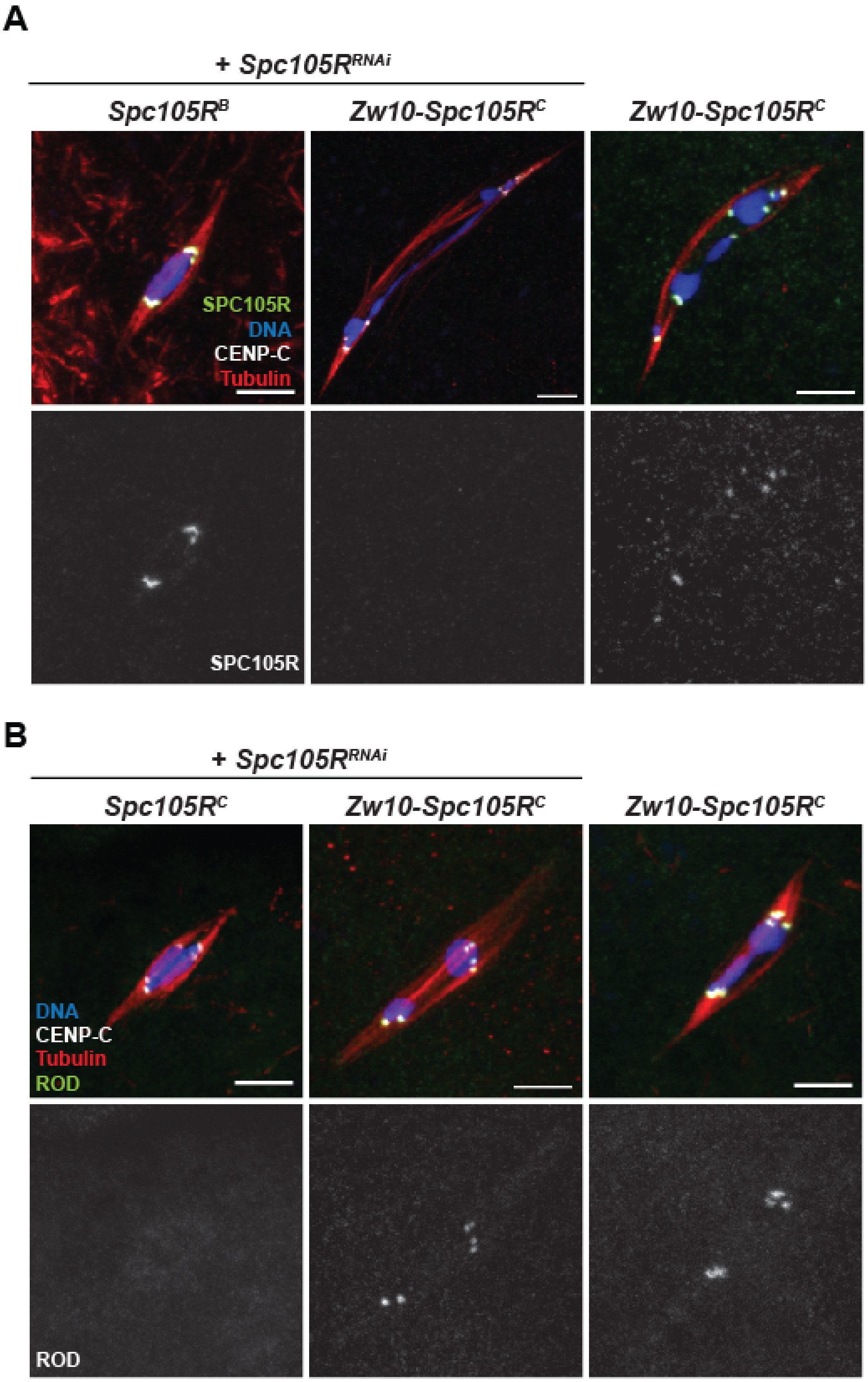
A
+ Spc105R^RNAi
Spc105R^B
Zw10-Spc105R^C
Zw10-Spc105R^C
SPC105R
DNA
CENP-C
Tubulin
SPC105R
B
+ Spc105R^RNAi
Spc105R^C
Zw10-Spc105R^C
Zw10-Spc105R^C
DNA
CENP-C
Tubulin
ROD
ROD

Figure S 5. **ROD and SPC105R localization in Z*w10-Spc105R*$^{C}$ oocytes.**

(A) SPC105R localization in Z*w10-Spc105R*$^{C}$, either in the presence or absence of *Spc105R*$^{RNAi}$. SPC105R (green) was detected using an antibody which recognizes the N-terminal regions of SPC105R and does not detect SPC105R$^{C}$. DNA is in blue, CENP-C in white, and tubulin in red. All images are maximum intensity projections of z stacks. (B) ROD$^{GFP}$ localization in oocytes expressing Z*w10-Spc105R*$^{C}$, either in the presence or absence of *Spc105R*$^{RNAi}$. ROD$^{GFP}$ is in green, DNA in blue, CENP-C in white, and tubulin in red. Scale bars represent 5 µm.

Figure S6

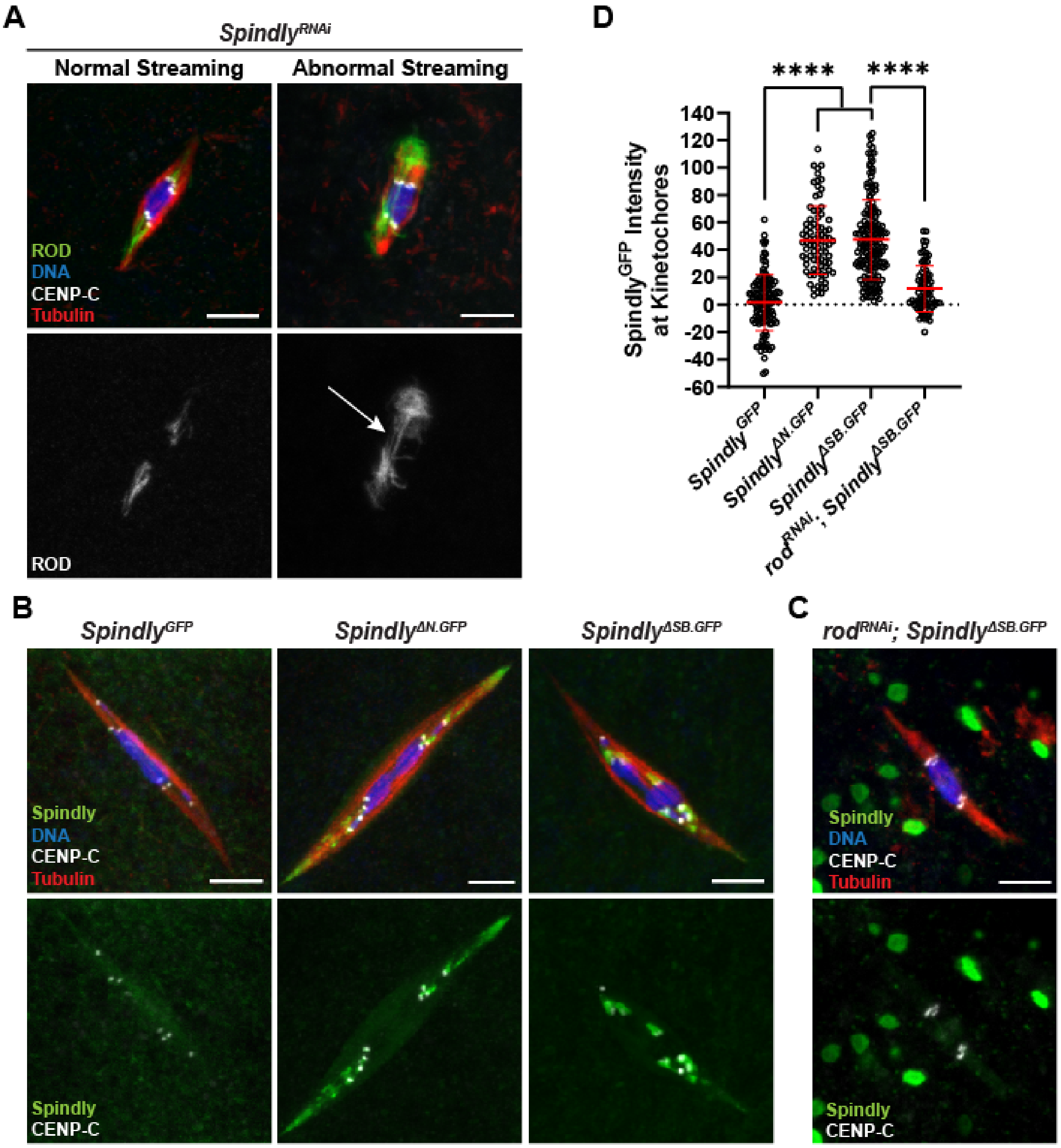
A
Spindly^RNAi
Normal Streaming
Abnormal Streaming
ROD
DNA
CENP-C
Tubulin
ROD
D
****
****
Spindly^GFP Intensity at Kinetochores
140
120
100
80
60
40
20
0
-20
-40
-60
Spindly^GFP
Spindly^ΔN.GFP
Spindly^ΔSB.GFP
rod^RNAi; Spindly^ΔSB.GFP
B
Spindly^GFP
Spindly^ΔN.GFP
Spindly^ΔSB.GFP
Spindly
DNA
CENP-C
Tubulin
Spindly
CENP-C
C
rod^RNAi; Spindly^ΔSB.GFP
Spindly
DNA
CENP-C
Tubulin
Spindly
CENP-C

Figure S 6. **Localization of Spindly variants and dependence of Spindly on ROD.**

(A) Examples of normal (left) and abnormal (right) $ROD^{GFP}$ streaming in *$Spindly^{RNAi}$* oocytes, with $ROD^{GFP}$ in green, DNA in blue, CENP-C in white, and tubulin in red. Single channel images (bottom) show $ROD^{GFP}$. Arrow points to abnormal streaming, which is defined as having $ROD^{GFP}$ in the central region between the centromeres. (B) $Spindly^{GFP}$ localization in wild-type or mutants of *Spindly*. $Spindly^{GFP}$ is in green, DNA in blue, CENP-C in white, and tubulin in red. (C) Localization of $Spindly^{\Delta SB.GFP}$ in *$rod^{RNAi}$* oocytes, with $Spindly^{GFP}$ in green, DNA in blue, CENP-C in white, and tubulin in red. $Spindly^{GFP}$ and CENP-C are shown below the merged images. All images are maximum intensity projections of z stacks. Scale bars represent 5 μm. (D) Quantification of $Spindly^{GFP}$ intensity at kinetochores in the indicated oocytes (from left to right, n = 123, 164, 83, and 75 kinetochores). Error bars show mean ± s.d.; ****$P<0.0001$ (unpaired two-tailed t test).
